# Quantifying (non)parallelism of gut microbial community change using multivariate vector analysis

**DOI:** 10.1101/2022.09.23.509066

**Authors:** Andreas Härer, Diana J. Rennison

## Abstract

Parallel evolution of phenotypic traits is regarded as strong evidence for natural selection and has been studied extensively in a variety of taxa. However, we have limited knowledge of whether parallel evolution of host organisms is accompanied by parallel changes of their associated microbial communities (i.e., microbiotas), which are crucial for their hosts’ ecology and evolution. Determining the extent of microbiota parallelism in nature can improve our ability to identify the factors that are associated with (putatively adaptive) shifts in microbial communities. While it has been emphasized that (non)parallel evolution is better considered as a quantitative continuum rather than a binary phenomenon, quantitative approaches have rarely been used to study microbiota parallelism. We advocate using multivariate vector analysis (i.e., phenotypic change vector analysis) to quantify direction and magnitude of microbiota changes and discuss the applicability of this approach for studying parallelism. We exemplify its use by reanalyzing gut microbiota data from multiple fish species that exhibit parallel shifts in trophic ecology. This approach provides an analytical framework for quantitative comparisons across host lineages, thereby providing the potential to advance our capacity to predict microbiota changes. Hence, we emphasize that the development and application of quantitative measures, such as multivariate vector analysis, should be further explored in microbiota research in order to better understand the role of microbiota dynamics during their hosts’ adaptive evolution, particularly in settings of parallel evolution.

## Introduction

Parallel evolution, the repeated evolution of similar traits in independent lineages in response to similar selective pressures, is a widespread phenomenon and provides strong evidence for natural selection (Colosimo et al., 2005, Steiner et al., 2009, Losos et al., 1998, Rosenblum et al., 2017, Elmer et al., 2010). Yet, the extent of parallelism varies considerably across levels of biological organization (e.g., genotype vs. phenotype) and across taxa (Bolnick et al., 2018). Substantial variation in parallelism can even be found among closely related populations adapting to seemingly similar habitats (Stuart et al., 2017). Traditionally, (non)parallel evolution has been regarded, and classified, as a binary phenomenon (evolution is parallel or not). However, it has recently been argued that by considering parallel evolution as a quantitative continuum, we will be better able to identify and understand the genetic and ecological factors that affect the extent of parallelism (Bolnick et al., 2018).

The study of parallelism has recently been extended to host-associated microbial communities and in particular the gut microbiota, the microbial community inhabiting a host’s gut (Ley et al., 2008b). To investigate microbiota parallelism, it can be useful to adopt both theoretical and methodological approaches developed for studying parallel evolution (e.g., Rennison et al., 2019). Gut microbial communities are highly diverse (Human Microbiome Project Consortium, 2012, Brooks et al., 2016, Youngblut et al., 2019) and affect host physiology in many ways (e.g., nutrient metabolism; Turnbaugh et al., 2006). The gut microbiota of an increasing number of host species is being characterized, and we are obtaining a more comprehensive picture of the extensive diversity of host-associated microbes (Song et al., 2020, Tarnecki et al., 2017). The gut microbiota is shaped by host genetics and ecological factors (Benson et al., 2010, Goodrich et al., 2014, Li et al., 2017, Sullam et al., 2012, Spor et al., 2011), and can impact the ecology and evolution of their hosts (Zepeda Mendoza et al., 2018, Rudman et al., 2019). Study systems in which closely related populations or species have independently adapted to similar ecological niches are particularly useful for studying the evolutionary ecology of host-associated microbial communities (e.g., Härer et al., 2020). In these systems, one can ask whether phenotypic or ecological changes that have occurred repeatedly in multiple host populations (i.e., parallel evolution), are associated with parallel changes in microbial communities (i.e., microbiota parallelism). We would like to emphasize that microbiota parallelism solely describes repeatability in the *direction* and *magnitude* of change of microbial communities, but not necessarily their parallel evolution. There is now growing interest in determining whether parallel adaptation of hosts is associated with parallel microbiota changes, as parallelism among independent gut microbial communities suggests changes could be predictable and adaptive (Delsuc et al., 2014, Song et al., 2020, Härer et al., 2020). Integration of microbiota data from a range of host populations that have repeatedly and independently adapted to similar ecological niches (e.g., Song et al., 2020) provides a powerful opportunity to investigate the ecological and evolutionary dynamics of host-microbe interactions.

Gut microbiota parallelism is predicted if hosts are adapting to similar trophic niches, since diet is known to be a major factor shaping gut microbial communities (Turnbaugh et al., 2009, Smits et al., 2017, Bolnick et al., 2014a). However, several ecological and genetic factors could further promote or hinder parallelism; these include similarity of host ecology, physiology and genetics, as well as, differential environmental exposure, and mode of microbial transmission (see Discussion for more details). This begs the question: Does the gut microbiota change in a predictable manner during their hosts’s parallel adaptation to similar trophic niches, and if so, what factors affect the likelihood of observing parallelism? To address questions of gut microbiota predictability and parallelism, we need hypothesis-driven tests leveraged in systems with well-characterized host ecology and repeated patterns of niche shifts. It is also imperative to employ quantitative statistical metrics. Here we suggest that multivariate vector analysis, a quantitative method, can be used to estimate the degree of microbiota parallelism, which might allow identifying the key ecological and evolutionary processes shaping variation in microbial communities. This method, originally termed ‘phenotypic change vector analysis’ was developed by Collyer and Adams for studying magnitude and direction of multivariate phenotypic change (Adams and Collyer, 2009, Collyer and Adams, 2007), and it has previously been applied to study variation in phenotypic (Stuart et al., 2017) and gut microbiota (Rennison et al., 2019) parallelism in threespine stickleback fish. In this method, vectors connecting the multivariate means (centroids) of microbial communities are estimated for pairs of host populations. The resulting vectors are then compared in a pairwise fashion for all host populations (figure 1; see Materials and Methods for more details). This approach provides information not only on the direction (angle between vectors), but also the magnitude (vector length) of microbiota divergence. However, it is the angle that quantifies parallelism, the smaller the angle between two population pairs, the more parallel the pattern of divergence (figure 1) (Stuart et al., 2017, Bolnick et al., 2018). Crucially, when integrated with additional ecological or genetic data, this quantitative approach allows direct tests of factors that affect direction and magnitude of microbiota changes.

**Figure 1:**
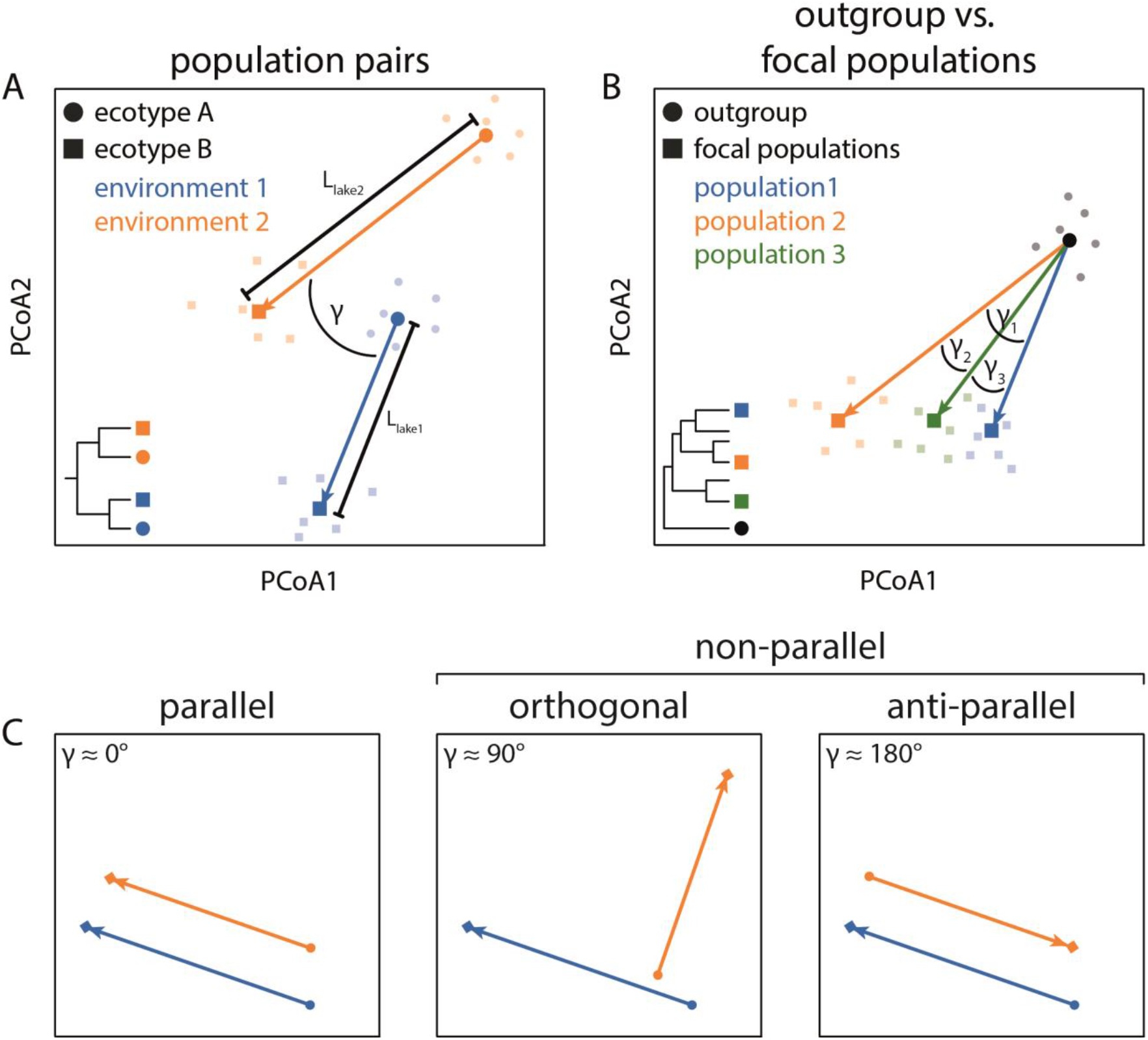
Illustration of vector analysis for determination of gut microbiota parallelism. Vectors connect the population means (centroids) between population pairs (A) or between an outgroup and several focal populations (B). The phylogenetic tree in (A) and (B) represent schematics to provide general information on the phylogenetic relationships of the studied populations. For the population pair comparisons, the two populations used to calculate a vector (γ) were most closely related to each other providing repeated and independent cases of ecological divergence. The populations or species included for each analysis are listed in table A1. Angles between vectors provide a quantitative measure of parallelism (C) and range from anti-parallel, to orthogonal to parallel (adopted from Bolnick et al., 2018). Angles between multivariate vectors were measured based on PCoA scores and represent a quantitative measure of gut microbiota parallelism. Vector lengths (L) provide information on the magnitude of gut microbiota changes. Centroids are shown as bold symbols (squares and circles) whereas individual data points are shown as faint symbols to illustrate differences in data distribution. Note that the direction of vectors is important and should be consistent across comparisons to obtain biologically meaningful results (e.g., always from ecotype A to ecotype B within a study system).

Parallel divergence in trophic ecology is well-documented in several teleost fishes; e.g., threespine stickleback (Bell and Foster, 1994, Taylor and McPhail, 1999), African and Neotropical cichlids (Elmer et al., 2014, Muschick et al., 2012), lake whitefish (Bernatchez et al., 1999) and Trinidadian guppies (Reznick et al., 1996) (table A1). To exemplify the utility of multivariate vector analysis for quantifying parallelism in compositional and functional changes of gut microbial communities, we reanalyzed published 16S rRNA gene sequencing data sets from these model systems. We discuss how estimates of magnitude of divergence and parallelism can be used to identify factors that affect microbial communities associated with many host lineages and give recommendations on the use of this approach. We further acknowledge current limitations of using multivariate vector analysis for studying microbiota parallelism and emphasize the need for further development of this approach in microbiota research. When applied to a wide range of host organisms, we argue that multivariate vector analysis has the potential to give a unique insight into the microbiota dynamics associated with their hosts’ adaptation to different ecological niches.

## Materials and Methods

### Data acquisition

We obtained 16S rRNA gene sequencing data from six published studies of parallel evolution in teleost fishes: herbivorous and carnivorous African cichlids (Baldo et al., 2017), benthic and limnetic Neotropical Midas cichlids (Härer et al., 2020), benthic and limnetic threespine stickleback from British Columbia (Rennison et al., 2019), freshwater and estuarine threespine stickleback from Oregon (Steury et al., 2019), benthic and limnetic lake whitefish (Sevellec et al., 2018), and low-predation and high-predation Trinidadian guppies (Sullam et al., 2015). Sample sizes for each population are indicated in table A1 and information on sequencing platform, amplicon region and NCBI archiving are listed in table A2. For each data set, all samples were included in a single sequencing run. To improve readability, we will refer to different host lineages as populations, whether they are populations of the same species or distinct species. We tested for gut microbiota parallelism (*i*) among population pairs, and (*ii*) between an outgroup and several focal populations (figure 1). Outgroups were selected based on phylogenetic information (see table A1). Marine threespine stickleback colonized freshwater environments around 10,000 – 12,000 years ago (Bell and Foster, 1994). Hence, a marine population was selected as the outgroup, since it represents the ancestral state. For Midas cichlids, the species *A. citrinellus* from Lake Nicaragua represents the outgroup to all crater lake species investigated, since crater lakes were colonized from the two great lakes of Nicaragua (L. Managua and L. Nicaragua) within the last 5,000 years (Kautt et al., 2020). In African cichlids, we selected a species from Barombi Mbo as the outgroup to several species from Lake Tanganyika based on a recent phylogeny by Irisarri et al. (2018). In this study system, the focal carnivorous populations were chosen to avoid phylogenetic clustering; the closest relatives of all focal populations differ in trophic ecology (e.g., herbivores or omnivores) (Baldo et al., 2017). Further information on the populations used can be found in table A1. Sequence data were downloaded from the NCBI Sequence Read Archive (SRA); information on sequencing platforms, sample sizes and accession numbers are provided in tables A1 and A2. Data was converted from SRA to FASTQ format using the fastq-dump function of the SRA Toolkit v2.9.6-1 (https://ncbi.github.io/sra-tools/).

### Gut microbiota analysis

Forward reads had higher sequence quality than reverse reads, and read lengths varied across studies due to differences in sequencing technology, which led to non-overlap of reads for some studies. Hence, we only used forward reads to achieve higher consistency in analysis across the different data sets. Forward reads were imported into the open-source bioinformatics pipeline QIIME2 (Bolyen et al., 2019).Sequence quality control was done with the plugin DADA2 (Callahan et al., 2016) and a phylogenetic tree was produced with FastTree 2.1.3 (Price et al., 2010). Taxonomy was assigned against the 16S rRNA gene Silva database version 132 (Quast et al., 2013) using the feature-classifier classify-sklearn plug-in in QIIME2 (Pedregosa et al., 2011). Taxonomic assignment was not done for Trinidadian guppies (Sullam et al., 2015) as this study used a different region of the 16S rRNA gene (V1-V3). Rarefaction depths and sequence lengths varied across data sets (table A1). We calculated different phylogenetic (weighted and unweighted UniFrac) and non-phylogenetic (Bray-Curtis dissimilarity) metrics for bacterial community composition (Lozupone et al., 2011). To infer metagenome function, MetaCyc pathway abundances, Kyoto Encyclopedia of Genes and Genomes (KEGG) orthologs and Enzyme Commission numbers were predicted with the PICRUSt2 plugin in QIIME2 (Douglas et al., 2020, Kanehisa et al., 2012) with a maximum nearest-sequenced taxon index (NSTI) cutoff of 2. Across the study systems, more than 90% of ASVs (92.7-99.5%) were below this cutoff, except for the study on whitefish where the proportion was slightly lower (85.3%; table A3). Mean and median NSTI scores ranged from 0.45-2.689 and 0.072-0.366, respectively. Based on distance matrices for all these different metrics, principal coordinate analyses (PCoA) were performed and PCoA scores were used as input for multivariate vector analyses.

### Multivariate vector analysis

We quantified compositional and functional gut microbiota parallelism using multivariate vector analysis. We largely followed the methodology reported in Rennison et al. (2019), which to date is the only study that has used this approach for studying gut microbiota parallelism to date. We found different degrees of parallelism for gut microbiota function than reported in the original study. This is likely due to differences in data processing and analysis pipelines, emphasizing the need to standardize data analysis when making inferences across studies. Multivariate vectors were calculated by connecting the population means (centroids) of PCoA scores, either between population pairs or between an outgroup and focal populations (as depicted in figure 1). The dimensionality of the data sets used to estimate angles for gut microbiota composition and function is listed in tables A4 and A5, respectively. Angles were measured between these vectors, and were calculated for all possible pairwise comparisons in each data set (e.g., between all three benthic-limnetic population pairs in threespine stickleback from British Columbia, Canada). The direction of vectors was held consistent, e.g., from benthic to limnetic across all comparisons within a study system, representing a repeated measure of evolutionary divergence between populations. Yet, we would like to mention that this does not necessarily reflect the direction of evolutionary change in all of our study systems (i.e., ancestral to derived). Angles for gut microbiota composition (Spearman’s ρ: 0.47 – 0.757) and function (Spearman’s ρ: 0.965 – 0.992) were highly reproducible across different diversity metrics (figures A1 & A2), and statistical tests for parallelism yielded largely similar results (tables A4 and A5). Hence, in the main text we only present Bray-Curtis dissimilarity for gut microbiota composition and MetaCyc pathway abundances based on Bray-Curtis dissimilarity for the inferred functional metagenome.

The angles provide us with a quantitative measure of parallelism within and across study systems. Smaller angles (below 90°) indicate parallelism, angles around 90° indicate orthogonal change and larger angles (above 90°) indicate anti-parallelism (figure 1C). A more detailed discussion on different interpretations of the distribution of angles, but also on the limitations of this method can be found in Bolnick et al. (2018) and in Watanabe (2022). To statistically test for gut microbiota parallelism, previous studies proposed to either regard changes as parallel when angles do not deviate from 0° (Bolnick et al., 2018) or when angles are significantly smaller than 90° (Rennison et al., 2019). To give us an initial simplistic indication of parallelism patterns, we used one-sample t-tests with an angle of 90° as the null expectation for non-parallelism since (almost) all data were normally distributed based on Shapiro-Wilk tests (Shapiro and Wilk, 1965). The only exception was gut microbiota composition of threespine stickleback population pairs from Rennison et al. (2019), for which we performed a one-sample Wilcoxon signed-rank test (Wilcoxon, 1945). However, due to the non-independence of pairwise angles (Watanabe, 2022), we also quantified parallelism by calculating distributions of random angles in multidimensional space (which is centered at 90°) and using Monte Carlo simulations (with 10^5^ iterations) to test for significant parallelism, or by performing a Rayleigh test which is used to examine the unimodal concentration of directional vectors (Mardia et al., 1979, Watanabe, 2022). We compare the results obtained using these different methods and discuss their interpretation.

To quantify the magnitude of gut microbiota changes, we calculated means of vector lengths (meanL) for each population pair. Correlation analyses were based on non-parametric Spearman rank correlation coefficients, as not all data were normally distributed. For the population pair comparisons, we also tested whether the taxonomic level at which microbial communities are studied affect the magnitude and direction of gut microbiota change based on the multivariate vector analysis. Taxonomic assignment was done in QIIME2 (more information provided above), biom tables were created for each study system at different taxonomic levels of the bacterial communities (phylum, class, order, family, genus, species). From each biom table, we produced a Bray-Curtis distance matrix and calculated PCoA scores. Estimates of angles and vector lengths were calculated based on these PCoA scores, similar to the other analyses mentioned above. All statistical analyses were done in R v3.5.1 (R_Core_Team, 2021).

## Results

### Gut microbiota parallelism across population pairs

First, we tested for parallelism across population pairs where vectors connect two closely related populations, representing cases of repeated divergence. Levels of gut microbiota parallelism varied considerably within and across teleost fish model systems for parallel evolution (figure 2). Within study systems, we detected a wide range of angles (e.g., 38.5 – 88.8° for gut microbiota composition in African cichlids; figure 2A), and parallelism estimates were also highly variable across study systems. When comparing mean angles against a null expectation of 90°, statistically significant parallelism was only found for gut microbial composition (mean: 68.89°; one-sample t-test: *P* < 0.001, *t* = -6.5658) and function (mean: 38.92°, *P* < 0.001, *t* = -12.513) among herbivorous and carnivorous African cichlids (figure 2A&B). We detected suggestive evidence for parallelism of gut microbial composition in benthic and limnetic threespine stickleback from British Columbia (mean: 81°, one sample Wilcoxon signed rank test: *P* = 0.087), but the sample size of this study was very small (n = 3). Similar results were obtained when using Monte Carlo simulations to compare mean angles against a multidimensional null distribution, we detected significant parallelism for African cichlids’ gut microbiota composition and function (*P* < 1 × 10^−5^ for both tests), and suggestive evidence in gut microbiota composition of benthic and limnetic threespine stickleback from British Columbia (*P* = 0.087) as well as in gut microbiota function of freshwater and estuarine threespine stickleback from Oregon (*P* = 0.052; figure A3). These findings were further supported by Rayleigh tests, in which significant concentrations of angles were detected for gut microbiota composition (S = 113.01, *P* < 1.26 × 10^−9^) and function (S = 160.47, *P* = 6.29 × 10^−21^) only in African cichlids. When performing multivariate vector analysis based on different taxonomic levels of gut microbial communities (species to phylum), we found that estimates of direction and magnitude were generally consistent, and independent of taxonomic resolution (figure 3).

**Figure 2:**
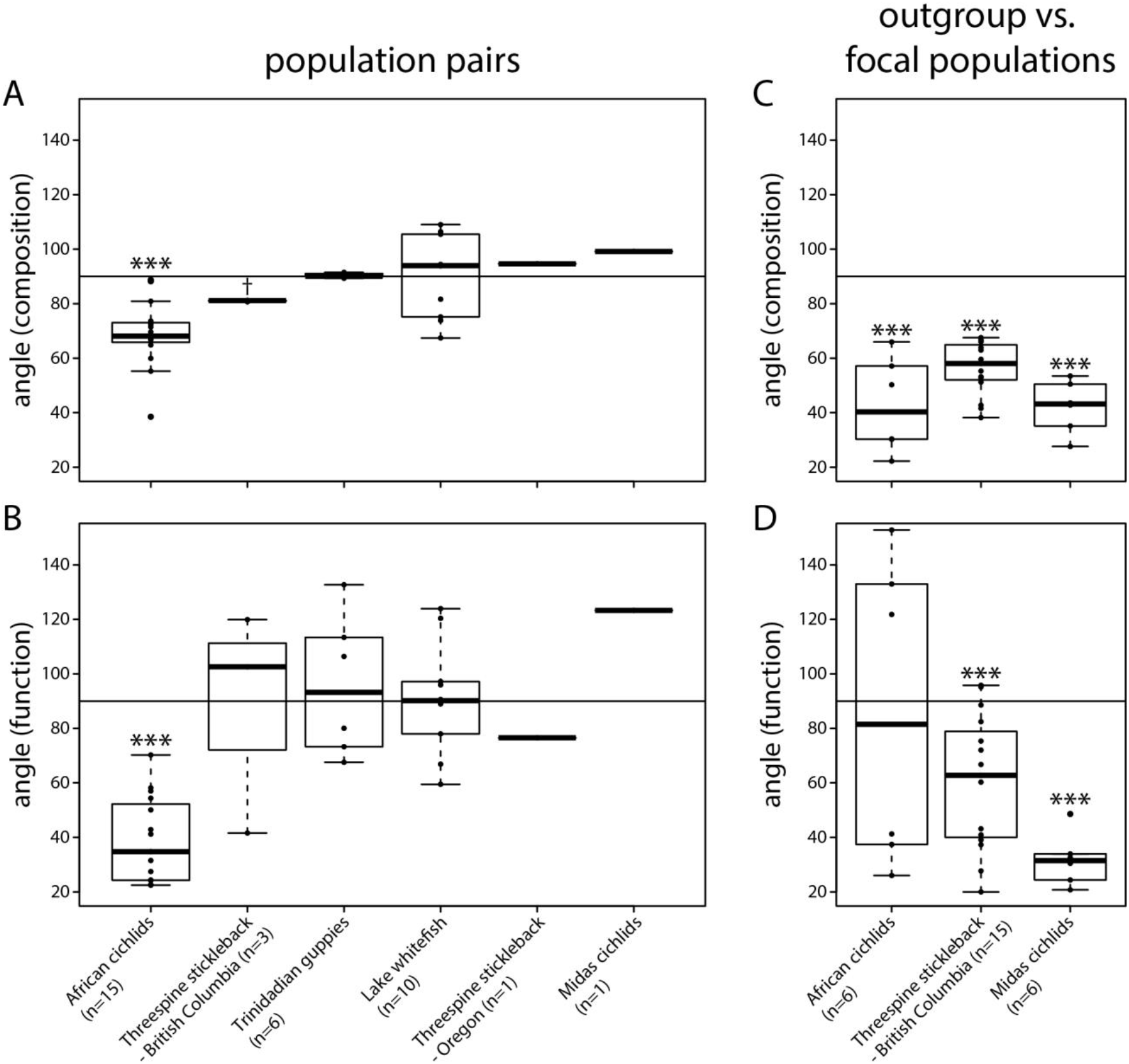
Significant gut microbiota parallelism among population pairs was detected in African cichlids for (A) gut microbiota composition and (B) function. In the outgroup comparisons, all three study systems showed significant gut microbiota parallelism for composition (C) and function (except for African cichlids; D). For all three study systems, outgroups inhabit distinct water bodies from the focal populations (table A1). Numbers of comparisons are indicated next to the name of each study system; the populations used for each analysis as well as samples sizes for each population are stated in table A1. Here, we show the results of testing mean angles against 90°, and no statistical tests were conducted for threespine stickleback population pairs from lakes and estuaries in Oregon and benthic and limnetic Midas cichlids from Nicaraguan crater lakes as we only had one comparison for each data set (A&B). †*P* < 0.1, ^*^*P* < 0.05, ^***^*P* < 0.001.

**Figure 3:**
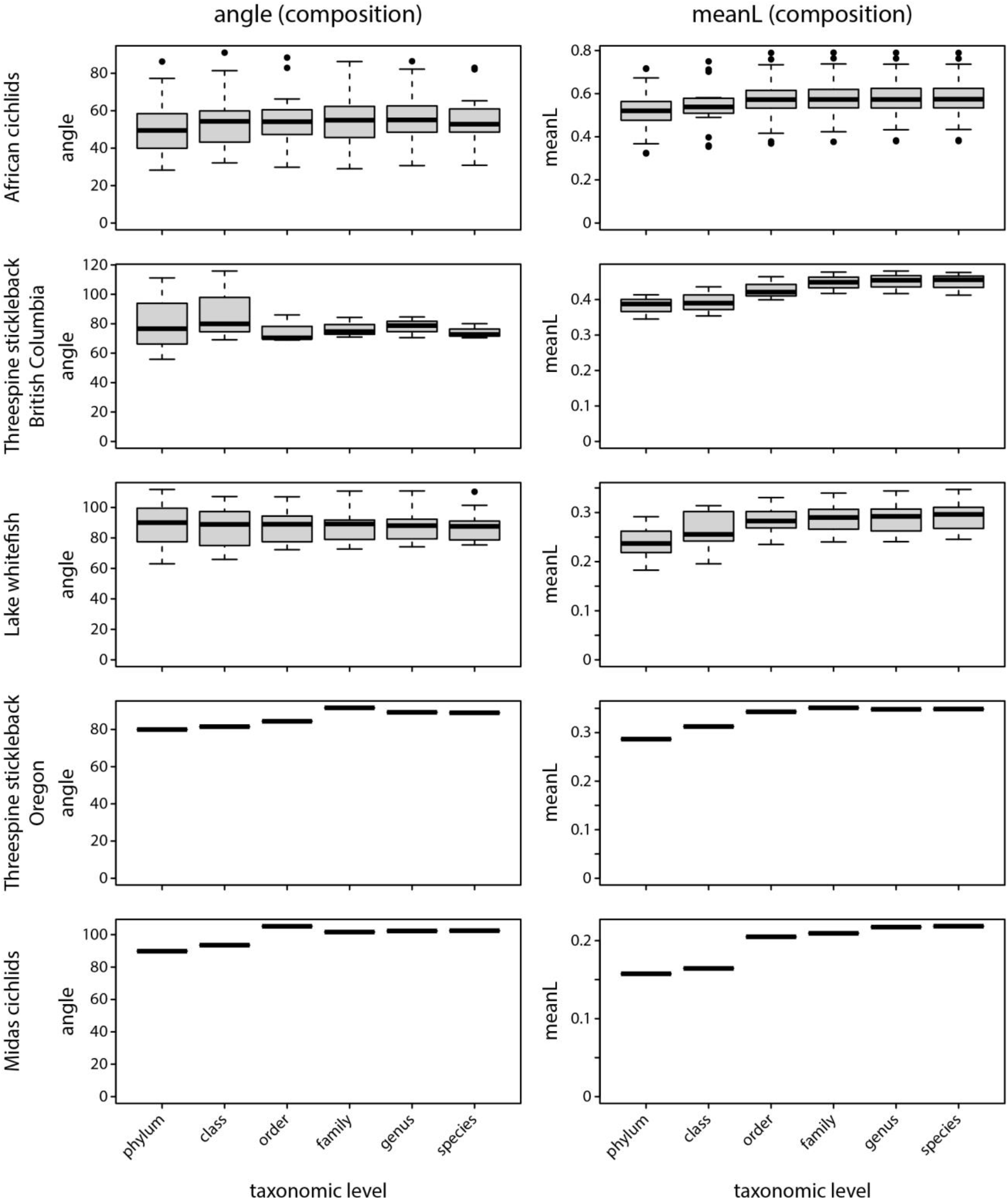
Estimates of direction (left column) and magnitude (right column) of gut microbiota change did not differ substantially with the taxonomic level used for multivariate vector analysis.

Angles for gut microbiota composition and function were strongly correlated for population pairs across study systems (Spearman’s ρ = 0.766, *P* < 0.001; figure 4A). On average, angles for gut microbiota function (mean: 80.9°) were smaller than for composition (mean: 70.3°), although this was not statistically significant (*P* = 0.096, *t* = 1.6987, figure 4B). When considering only comparisons that showed evidence of gut microbiota parallelism (i.e., angles were smaller than 90°), gut microbiota function angles were significantly smaller (i.e., more parallel) (*P* < 0.001, *t* = 5.5881, figure A4). There was also a significant negative correlation between gut microbiota parallelism (angles) and the mean magnitude of gut microbial change (meanL) for composition (ρ = -0.706, *P* < 0.001) and function (ρ = -0.581, *P* < 0.001; figure A5A&B).

**Figure 4:**
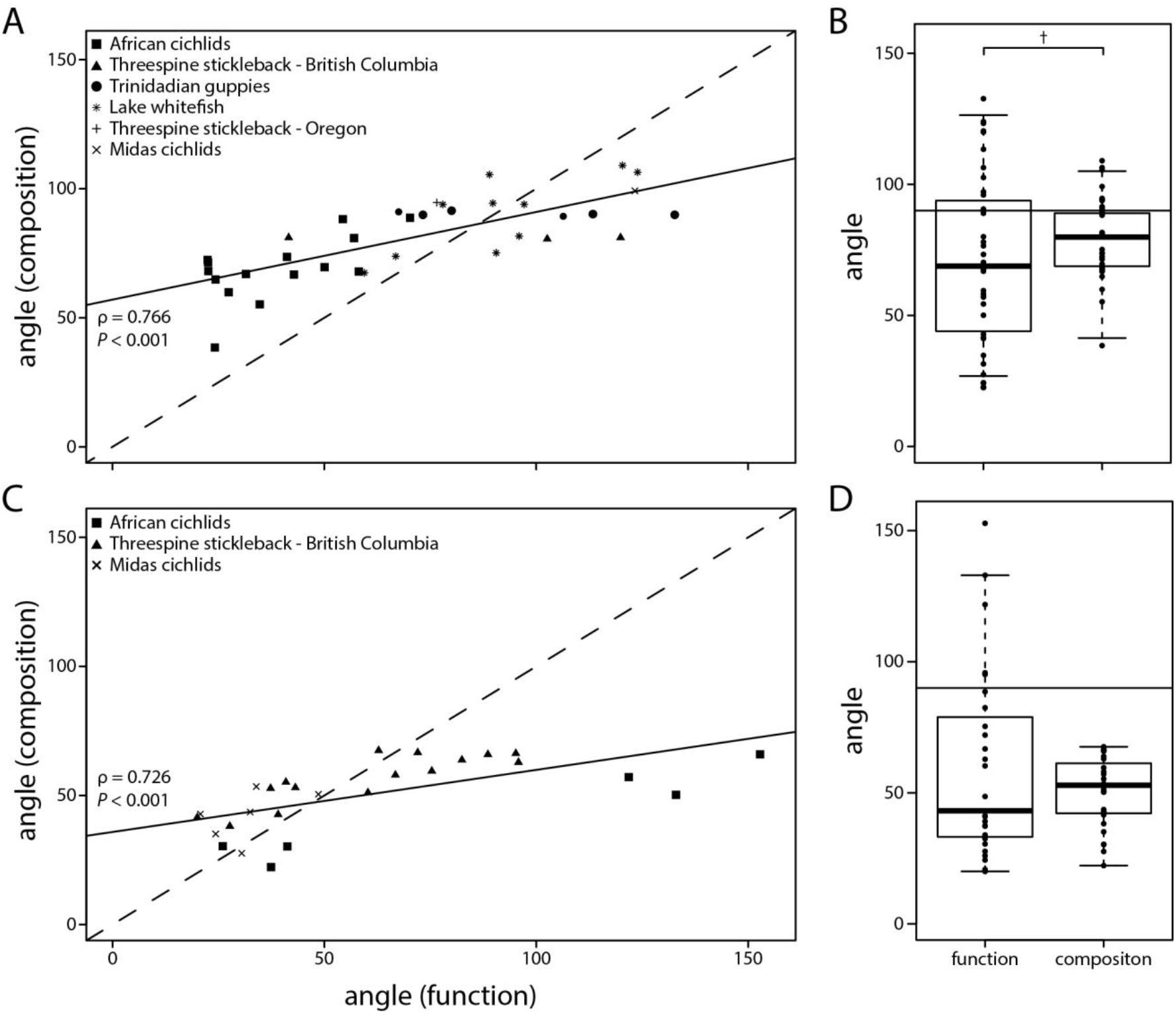
Levels of gut microbiota (non)parallelism were strongly correlated for gut microbiota composition and function for population pairs (A) and the outgroup comparisons (C); the dashed line has a slope of 1. There was only suggestive evidence for a difference in angles between gut microbiota composition and function for population pairs (B) but not for the outgroup comparison (D). Information

### Gut microbiota parallelism across focal populations compared to outgroup

When comparing an outgroup to focal populations adapted to similar ecological niches (figure 1B), we found strong evidence for compositional and functional gut microbiota parallelism across all three study systems; almost all angles were smaller than 90° (figure 2C&D).

By comparing mean angles against a null expectation of 90°, gut microbiota changes associated with carnivory in African cichlids from Lake Tanganyika (compared to an herbivorous outgroup from Lake Barombi Mbo) were significantly parallel for composition (mean: 42.7°, *P* < 0.001, *t* = -6.6208), but not for function (mean: 85.4°, *P* = 0.425, *t* = -0.2006). Gut microbiota changes in freshwater benthic and limnetic threespine stickleback ecotypes from British Columbia were significantly parallel when compared to the ancestral marine population (composition, mean: 56.4°, *P* < 0.001, *t* = -13.378; function, mean: 60.5°, *P* < 0.001, *t* = -4.6348). In benthic and limnetic Midas cichlids from two crater lakes, we also detected significant gut microbiota parallelism compared to the ancestral population from great lake Nicaragua (composition, mean: 42.2°, *P* < 0.001, *t* = -12.208; function, mean: 31.8°, *P* < 0.001, *t* = -14.772). We obtained similar results when using Monte Carlo simulations to compare mean angles against a multidimensional null distribution. Angles were significantly parallel for gut microbiota composition (*P* < 1 × 10^−5^) but not for function (*P* = 0.25) in African cichlids. In threespine stickleback from British Columbia, we detected significant parallelism for composition and function (*P* < 1 × 10^−5^ for both tests), the same was true for benthic and limnetic Midas cichlids (*P* < 1 × 10^−5^ for both tests; figure A6). Using Rayleigh tests, we detected significant concentrations of angles for gut microbiota composition (S = 59.53, *P* < 6.29 × 10^−7^) but not for function (S = 14.92, *P* = 0.25) in African cichlids. In threespine stickleback from British Columbia, angles were significantly concentrated for composition (S = 133.77, *P* < 4.64 × 10^−15^) and function (S = 92.39, *P* < 2.81 × 10^−10^). The same held true for composition (S = 294.37, *P* < 2.32 × 10^−28^) and function (S = 248.02, *P* < 1.17 × 10^−26^) in benthic and limnetic Midas cichlids.

Angles for gut microbiota composition and function were strongly correlated across study systems (Spearman’s ρ = 0.726, *P* < 0.001; figure 4C). Angles did not differ between gut microbiota composition and function (*P* = 0.21, *t* = -1.2787; figure 4D); even when only considering comparisons with angles below 90° for both measures (*P* = 0.723, *t* = 0.3579). There was no correlation between gut microbiota parallelism (angles) and mean magnitude of gut microbial change (meanL), for composition (ρ = -0.08, *P* = 0.691) or function (ρ = 0.073, *P* = 0.716; figure A5C&D).

## Discussion

Multivariate vector analysis offers the opportunity to quantify direction and magnitude of microbiota changes, thereby allowing identification of factors that shape microbial communities across host populations. The purpose of our study is to exemplify and advocate the use of this approach in microbiota research, compare different statistical approaches to test for parallelism, discuss current limitations, and give recommendations on methodological aspects. To this end, we applied multivariate vector analysis to investigate the gut microbiota of several classic teleost fish model systems that exemplify parallel ecological shifts. While this manuscript is focused on the study of parallelism, multivariate vector analysis can also be used to study the extent of convergence or divergence of microbiota changes, when incorporating information on the direction of evolutionary change in host lineages (Bolnick et al., 2018). It should be noted that multivariate vector analysis has only been used in one microbiota study (Rennison et al., 2019), and more conceptual and methodological research is needed to determine how characteristics of microbiota data sets affect parallelism estimates in order to comprehensively interpret the biological implications of such results. We would also like to emphasize that the small set of study systems included here are meant to exemplify the use of this quantitative approach. The goal was not to identify or disentangle the effects of specific factors that shape gut microbiota parallelism, but we encourage future studies to apply quantitative analyses to larger data sets in order to investigate general patterns and processes using the methods presented here.

### Multivariate vector analysis to quantify microbiota parallelism

When studying microbiota parallelism, one admittedly simplified prediction is that parallel adaptation of hosts to a novel diet translates to parallel changes of their gut microbiota. However, previous studies using a diversity of statistical approaches suggested that parallel shifts of the gut microbiota are not necessarily expected or commonly observed. For example, across several model systems of parallel trophic divergence, gut microbiota parallelism has previously been reported only for African cichlids (Baldo et al., 2017) and benthic and limnetic threespine stickleback (Rennison et al., 2019). There has been no conclusive evidence presented to suggest gut microbiota parallelism in Nicaraguan Midas cichlids (Härer et al., 2020), lake whitefish (Sevellec et al., 2018) or Trinidadian guppies (Sullam et al., 2015).

The variation in gut microbiota parallelism among systems described in previous studies might be biologically real or could be due to differences in the methodological approaches used, which limits our ability to make inferences across systems. Different statistical methods were used to analyze the gut microbiota data of these species; four of the studies used permutational analysis of variance (PERMANOVA) to determine the effects of ecotype, the environment, and their interaction to infer parallelism. One study on African cichlids further inferred parallelism based on PCoA scores of the first axes (Baldo et al., 2017). A drawback of such approaches is that gut microbiota changes are scored as parallel or non-parallel in a binary manner, based on a given significance threshold. Such methods also do not provide estimates of the magnitude of change. Thus, we lack quantitative information on the extent and variation of (non)parallelism (figure 1C). Multivariate vector analysis allows quantification of variation across independent population pairs. When applied to large data sets, the method can facilitate identification of factors that underlie variation in parallelism. This is important as we know that a multitude of genetic and ecological factors affect the composition of microbial communities in different host lineages, including teleost fishes (Amato et al., 2019, Bletz et al., 2016, Burns et al., 2017, Zhang et al., 2016, Bolnick et al., 2014b, Baldo et al., 2017). When applied to our six case studies, we found similar results as other statistical methods; there was only strong evidence of parallelism among population pairs of African cichlids, and weaker evidence for benthic-limnetic threespine stickleback (figure 2A&B). The consistency of these results highlights the suitability of multivariate vector analysis for studying microbiota parallelism, with the added value of quantitative data that can be used in direct hypothesis testing.

### Methodological considerations

We tested whether methodological aspects, such as the choice of metric for determining differences in bacterial community composition (e.g., Bray-Curtis dissimilarity, weighted and unweighted UniFrac) or the taxonomic level at which microbial communities are studied, affect estimates of direction and magnitude of gut microbiota change. We detected strong correlations among angle estimates generated using the different metrics (figure A1), and statistical significance of parallelism was generally consistent across different metrics for both taxonomic composition (table A4) and inferred metagenome function (table A5). This suggests that the parallelism estimates from multivariate vector analysis are not strongly affected by the choice of diversity metric. It should be noted that multivariate vector analysis is not restricted to analyzing data from principal coordinates analysis (PCoA); data from other ordination methods, such as non-metric multidimensional scaling (NMDS), can also be used as input for the analysis. NMDS data have been used in a parallelism analysis of the threespine stickleback gut microbiota, and the results are qualitatively similar to the PCoA analysis reported here (Rennison et al., 2019). Further, results were robust across taxonomic levels (figure 3), which was a bit surprising as the composition of microbial communities might be more stochastic at lower taxonomic levels. Thus, predictability of microbiota change (i.e., parallelism) could have been expected to be stronger at higher taxonomic levels. However, our results suggest that researchers should be able to tailor the taxonomic level to the needs of their particular study.

Previous studies utilized different approaches for significance testing of angles from multivariate vector analysis (Bolnick et al., 2018, Rennison et al., 2019). One study that to infer parallelism, methods should be based on the distribution of random angles, which depends on data dimensionality (Watanabe, 2022), rather than testing against a certain angle (e.g., 90°). Fortunately, in highly multidimensional space, such as that of microbiota data, random angles approximately follow a normal distribution centered around 90° (Watanabe, 2022). Hence, we argue that one-sample t-tests comparing the mean angles of our empirical data against the null expectation of 90° is a useful first test of parallelism. However, given that the 90° value is only a rough approximation, we implemented two additional statistical analyses to test for significant parallelism. We used Monte Carlo simulations to compare the estimated mean empirical angles against a multidimensional null distribution. We further implemented Rayleigh tests which test for the unimodal concentration of directional vectors (Mardia et al., 1979). Across these three statistical methods, we obtained largely consistent results. Since each statistical approach tests for a different property (see Watanabe, 2022 for a more detailed discussion), the use of multiple approaches helps obtain a more comprehensive and robust picture of whether changes of the gut microbiota are consistent with a parallel, orthogonal or anti-parallel pattern of change (figure 1C).

Multivariate vector analysis was developed for studying a range of phenotypic traits (Adams and Collyer, 2009), and has been used to study morphological and behavioral parallelism (Stuart et al., 2017). Only one study has applied this approach for studying gut microbiota parallelism in threespine stickleback (Rennison et al., 2019), hence, many open questions remain considering the application of multivariate vector analysis in microbiota research. The highly diverse and dynamic nature of microbial communities (e.g., Smits et al., 2017, Youngblut et al., 2019) could affect the interpretation of parallelism estimates. For example, more work is needed to determine how microbiota dispersion within populations and overlap among host populations can affect parallelism estimates, as well as their biological implications. Simulations of microbiota data could be leveraged to quantify how variation in these factors affects the range of possible angles observed, which would improve the interpretation of parallelism estimates. At the same time, more empirical studies are needed to obtain a comprehensive picture of microbiota parallelism in a phylogenetically and geographically broad range of host lineages.

### Progress and prospects in implementing multivariate vector analysis

Multivariate vector analysis quantifies changes in microbial communities and enables testing the effects of ecological or genetic factors on parallelism. Across six model systems of parallel evolution, we observed extensive variation in gut microbiota parallelism (figure 2), and we sought to conduct preliminary analyses to explore some of the factors that might affect parallelism. For example, we found that changes in gut microbiota function might be more parallel than composition, which is in line with findings of individual studies (figure 4) (Rennison et al., 2019, Härer et al., 2020). This pattern could be explained by the functional redundancy within microbial communities (Ley et al., 2006). Variation in taxonomic composition might be produced by historical contingency, priority effects or microbial dispersal ability (Costello et al., 2012, Martinez et al., 2018). Yet, taxonomically distinct bacterial taxa can provide similar metabolic functions, potentially causing stronger signatures of parallelism in gut microbiota function when hosts adapt to similar ecological niches. At the same time, parallelism estimates for gut microbiota composition and function were strongly correlated (figure 4), indicating that similar factors shape the extent of parallelism. We suggest that work integrating parallelism estimates for both composition and function from a variety of disparate host taxa is needed to explore this pattern further. But, it is important to be cautious when interpreting results on inferred metagenome function using tools such as PICRUSt2, particularly in non-model organisms (Douglas et al., 2020); the reliability of metagenome prediction highly depends on the database of available bacterial genomes, which can vary across host organisms (Sun et al., 2020).

Shifts in gut microbial communities were found to be most parallel among population pairs when the overall magnitude of divergence (meanL) was greatest (figure A5); this raises the question of whether substantial divergence in the gut microbiota might indicate shared adaptive changes. Strong selection pressures may lead to more determinism in microbial community assembly and, consequently stronger microbiota parallelism. In contrast, when there is little microbiota divergence, differences might be produced mainly by stochastic processes, e.g., priority effects and drift (Martinez et al., 2018). These processes are unlikely to generate similar microbiota shifts, and theoretical work suggests that parallelism may only be seen when the selection landscape is highly parallel (Thompson et al., 2019). For morphological traits and genetic divergence, it has been shown that the degree of environmental variation among evolutionary replicates directly predicts the magnitude of parallelism based on multivariate vector analysis (Stuart et al., 2017). If more similar selection pressures translate to more parallel host phenotypic and ecological change, this could lead to similar shifts in microbial communities. Accordingly, host adaptation to similar diets, when accompanied by changes in gut morphology, is expected to promote gut microbiota parallelism (Muegge et al., 2011, Ley et al., 2008a, Baldo et al., 2017). Yet, strong parallelism could also be explained by variation in environmental factors, as seen in our outgroup comparisons (figure 2C&D). In these settings, focal populations and the outgroups live in strongly differentiated environments (Torres-Dowdall et al., 2017, Barluenga and Meyer, 2010, Ormond et al., 2011), suggesting that similar patterns of divergence (parallelism) might be primarily driven by abiotic (physicochemical properties) and biotic (microbial communities) differences between environments, in addition to or instead of host trophic ecology. This was further supported by findings in Midas cichlids and threespine stickleback, where angles did not appear to be smaller for comparisons that only included the outgroup and a certain ecotype (i.e., benthics or limnetics) than for comparisons that included the outgroup and both ecotypes (figure A7). Yet, one should be careful when drawing conclusions from these results since sample sizes were small. Future work that estimates the similarity of abiotic and biotic factors among host populations will be key to determining whether parallel selective regimes generally translate to parallelism in changes of microbial communities.

Variation in microbial communities can be strongly associated with host phylogeny and the extent of genetic divergence (Brooks et al., 2016, Youngblut et al., 2019, Benson et al., 2010, Goodrich et al., 2014, Li et al., 2017); thus, increasing genetic (and phylogenetic) distance among hosts (Smith et al., 2015) could affect the likelihood of observing microbiota parallelism. Stronger parallelism might be predicted for host lineages that split earlier, with sufficient time to diverge ecologically. Results from Neotropical and African cichlids hint at such an association; the very recently diverged crater lake Midas cichlids from Nicaragua (< 5,000 years ago) (Kautt et al., 2020) showed no evidence for gut microbiota parallelism. In contrast, there was strong evidence in much older African cichlid lineages, where divergence times are in the range of millions of years (figure 2) (Baldo et al., 2017). Threespine stickleback and whitefish colonized freshwater lakes after the last ice age (< 12,000 years ago) and formed ecologically distinct species pairs adapted to different niches (Schluter and McPhail, 1992, Matthews et al., 2010, Bernatchez et al., 1999). Yet, we only detected some evidence for parallelism in benthic-limnetic stickleback (figure 2), suggesting that an association with host divergence time is not necessarily expected in general. Parallelism was also stronger in the outgroup comparisons (figure 2), which could be driven by stronger genetic divergence of the outgroup compared to focal populations (but also by environmental differences, see previous paragraph). However, for two of the study systems (Midas cichlids and threespine stickleback) the outgroup and the focal populations split very recently (<12,000 years), whereas population pairs of African cichlids split much earlier. Hence, the stronger parallelism in the outgroup comparisons cannot be explained solely by host divergence time. Again, a diverse sampling of host taxa, potentially including intra-as well as interspecific host comparisons, will be key to robustly test whether divergence time is a key factor determining patterns of gut microbiota parallelism.

There are numerous possible uses of multivariate vector analysis to investigate changes in host-associated microbial communities as many genetic and ecological factors may determine parallelism. For example, it would be particularly interesting to compare patterns among trophic specialists and generalists. Specialists might have a more stable gut microbiota compared to the gut microbiota of generalists, which is expected to fluctuate more over time (Smits et al., 2017, Baniel et al., 2021). Thus, this variation in gut microbiota plasticity (Kolodny and Schulenburg, 2020) could affect observed parallelism. The mode of gut microbiota transmission could be another interesting factor to consider using multivariate vector analysis. The acquisition of gut microbiota from environmental sources (i.e., horizontally) vs. transfer from mother to child (i.e., vertically) could affect the likelihood of observing parallelism, depending on whether these sources are shared among hosts (Mulder et al., 2009, Smith et al., 2015). To obtain a comprehensive understanding of the evolutionary ecology of the gut microbiota, future studies should make an effort toward combining quantitative measures with a diverse range of genetic and phenotypic host data, as well as environmental data.

### Conclusions

Our study exemplifies the use of multivariate vector analysis for studying microbiota dynamics, and discusses potential advantages compared to more commonly used statistical approaches. By using a common analytical framework, multivariate vector analysis allows quantification of the direction and magnitude of microbiota change. When applied across a broad range of host taxa, these estimates can be leveraged to examine general patterns of repeatability. Combining quantitative estimates with host-associated and environmental data offers the possibility to improve our knowledge of the eco-evolutionary processes that shape microbial community dynamics. Identification of these factors will improve our ability to predict (putatively adaptive) shifts in microbial communities. Hence, we encourage further adoption of quantitative measures for studying microbiota dynamics during adaptive evolution of their hosts, particularly in settings of parallel evolution.

## Data Accessibility

All sequencing data included in this study has been obtained from data sets published in the NCBI Sequence Read Archive (SRA) and can be obtained under the following project numbers: PRJNA341982 (African cichlids), PRJNA615202 (Nicaraguan cichlids), PRJNA475955 (threespine stickleback from British Columbia), PRJNA657232 (threespine stickleback from Oregon), PRJNA394764 (lake whitefish), and PRJNA259592 (Trinidadian guppies). Further information on the populations used for our analyses, sequencing platforms, amplicons, and the associated accession numbers are stated in tables A1 and A2.

## Author Contributions

A.H. and D.J.R conceptualized the study, D.J.R. developed the vector analysis pipeline and scripts, A.H. analyzed the data using these resources. A.H. wrote the manuscript with input from D.J.R.

## Competing Interests

The authors declare no competing interests.

## Acknowledgments

This work was supported by funding from the Deutsche Forschungsgemeinschaft (DFG, German Research Foundation) – project number 458274593 to A.H. and from the University of California San Diego to D.J.R.

## Appendix A

**Figure A1:**
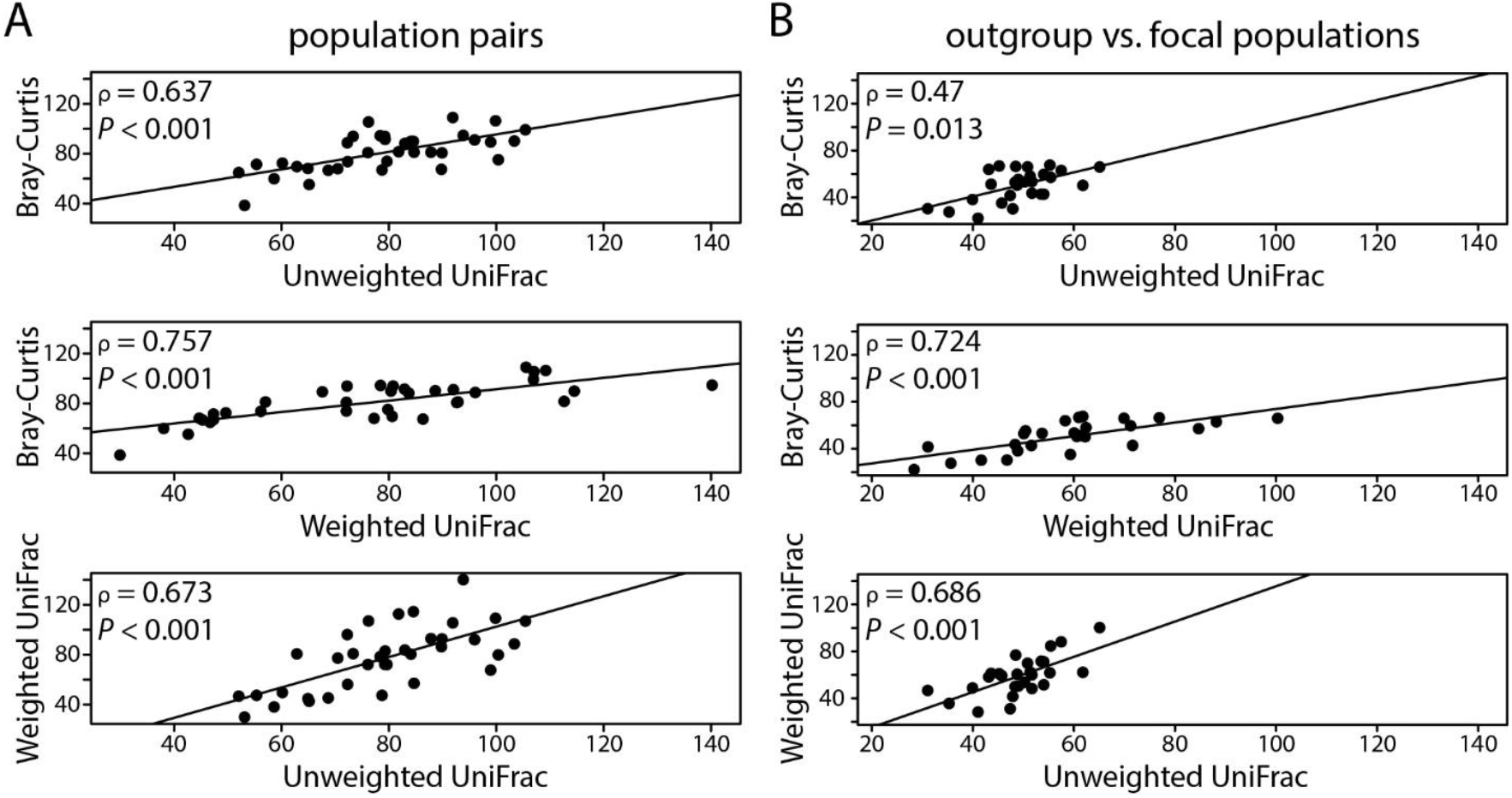
Parallelism estimates among different gut microbiota composition metrics were strongly correlated across comparisons of (A) population pairs and (B) outgroups vs. focal populations. Angles calculated from the multivariate vector analysis are indicated on the x- and y-axes.

**Figure A2:**
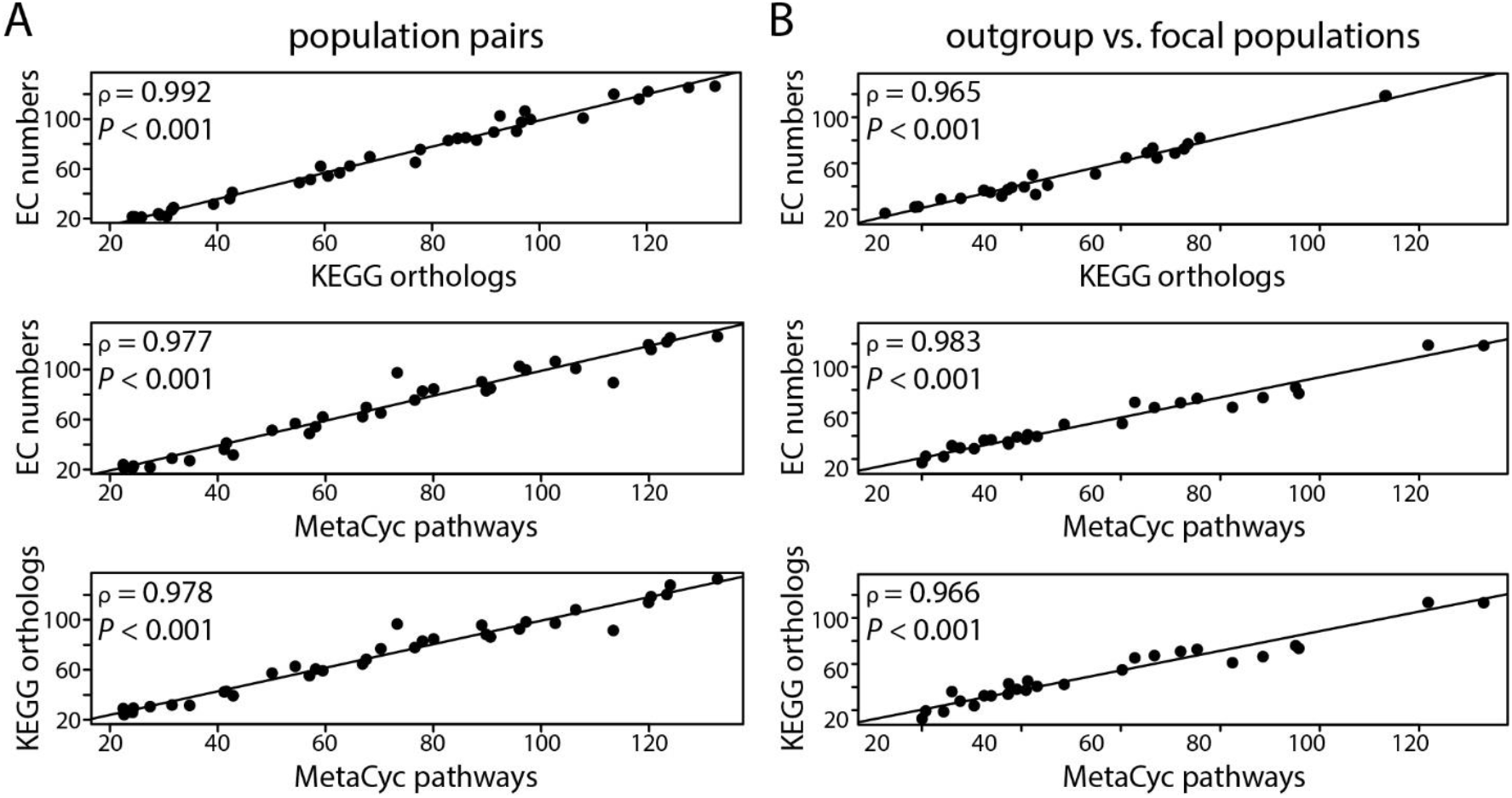
Parallelism estimates among different gut microbiota function metrics were strongly correlated across comparisons of (A) population pairs and (B) outgroups vs. focal populations. Angles calculated from the multivariate vector analysis are indicated on the x- and y-axes.

**Figure A3:**
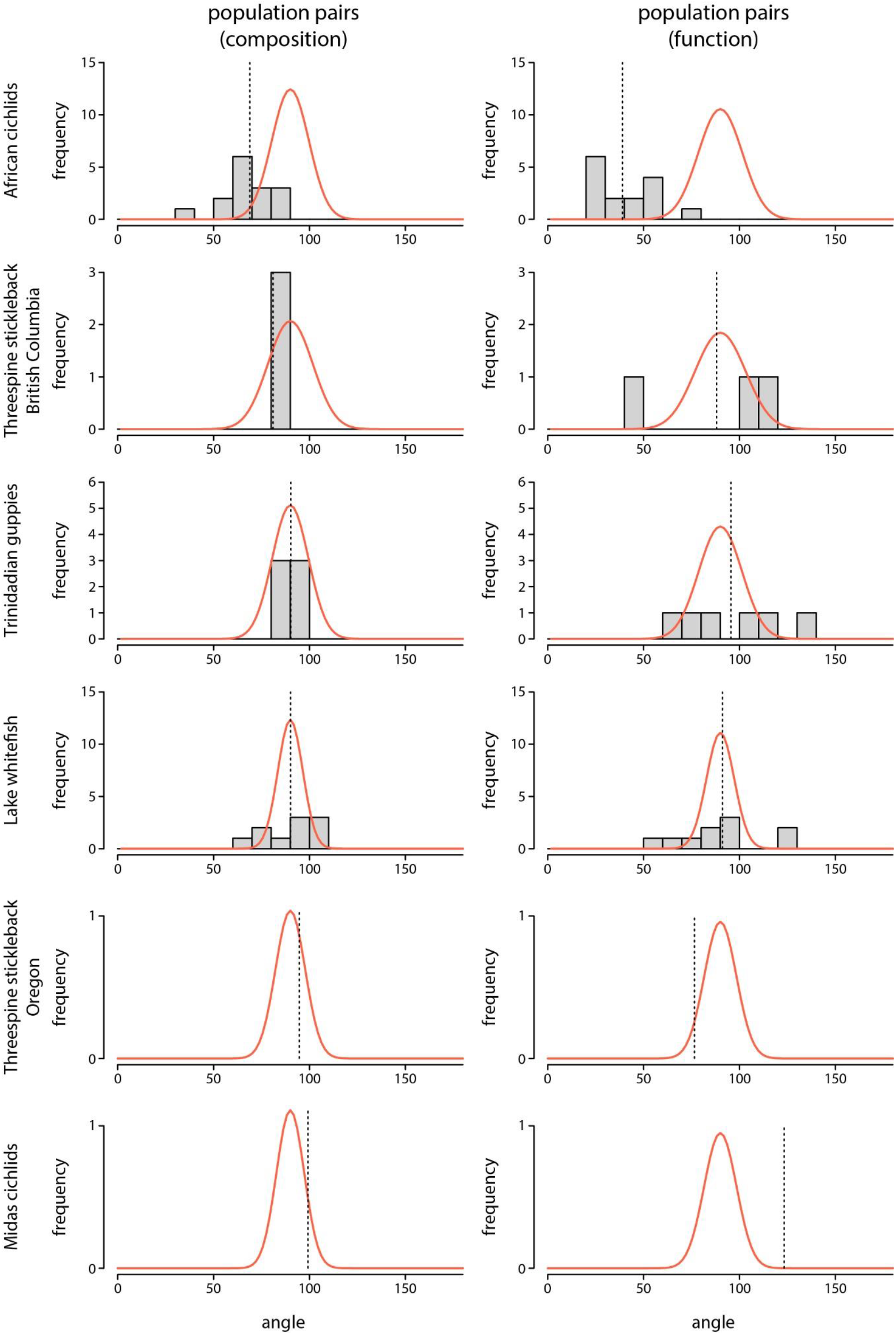
Histograms showing the distribution of angles among population pairs for gut microbiota composition (left column) and function (right column). The vertical dashed line represents the mean angle for each study system, and the multidimensional distribution of random angles is illustrated by the red curves, the shape of a curve is determined by the number of PCoA axes included in each analysis.

**Figure A4:**
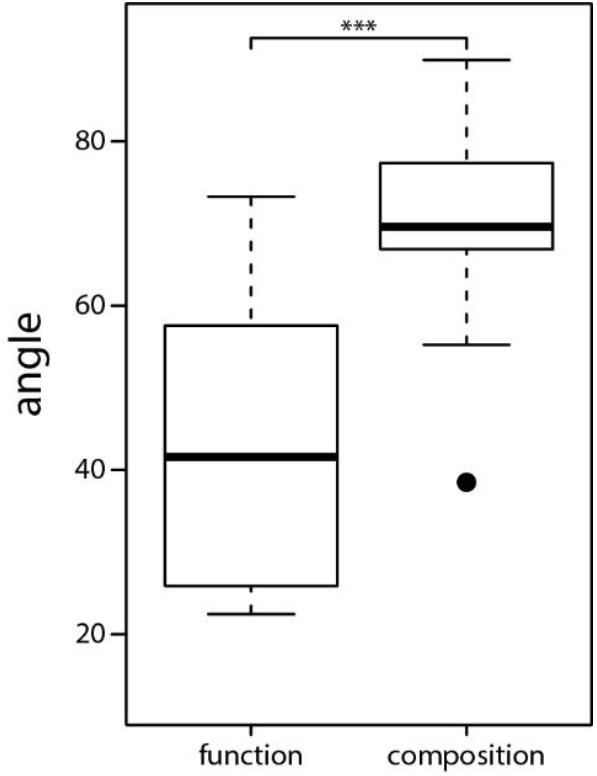
Across population pair comparisons, angles for gut microbiota function were significantly smaller than for composition (*P* < 0.001, *t* = 5.5881) when only considering comparisons that showed evidence of gut microbiota parallelism (angles < 90° for composition and function).

**Figure A5:**
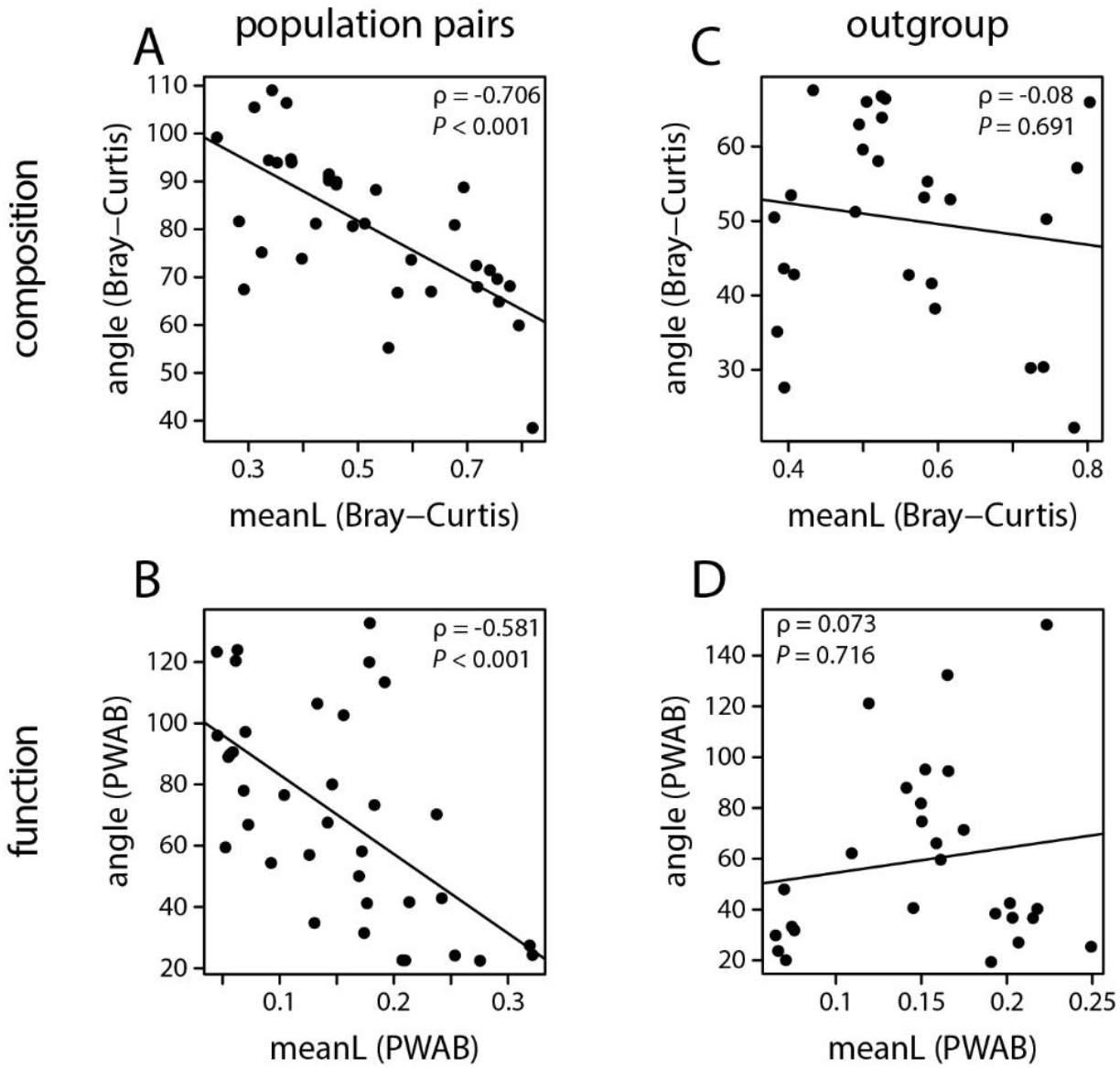
The strength of parallelism (angles) and the mean magnitude of gut microbiota change (meanL) were correlated for composition and function across population pair comparisons (A&B); the more parallel gut microbiota changes were (i.e., smaller angles), the higher the magnitude of change. No correlation was observed for outgroup vs. focal population comparisons (C&D).

**Figure A6:**
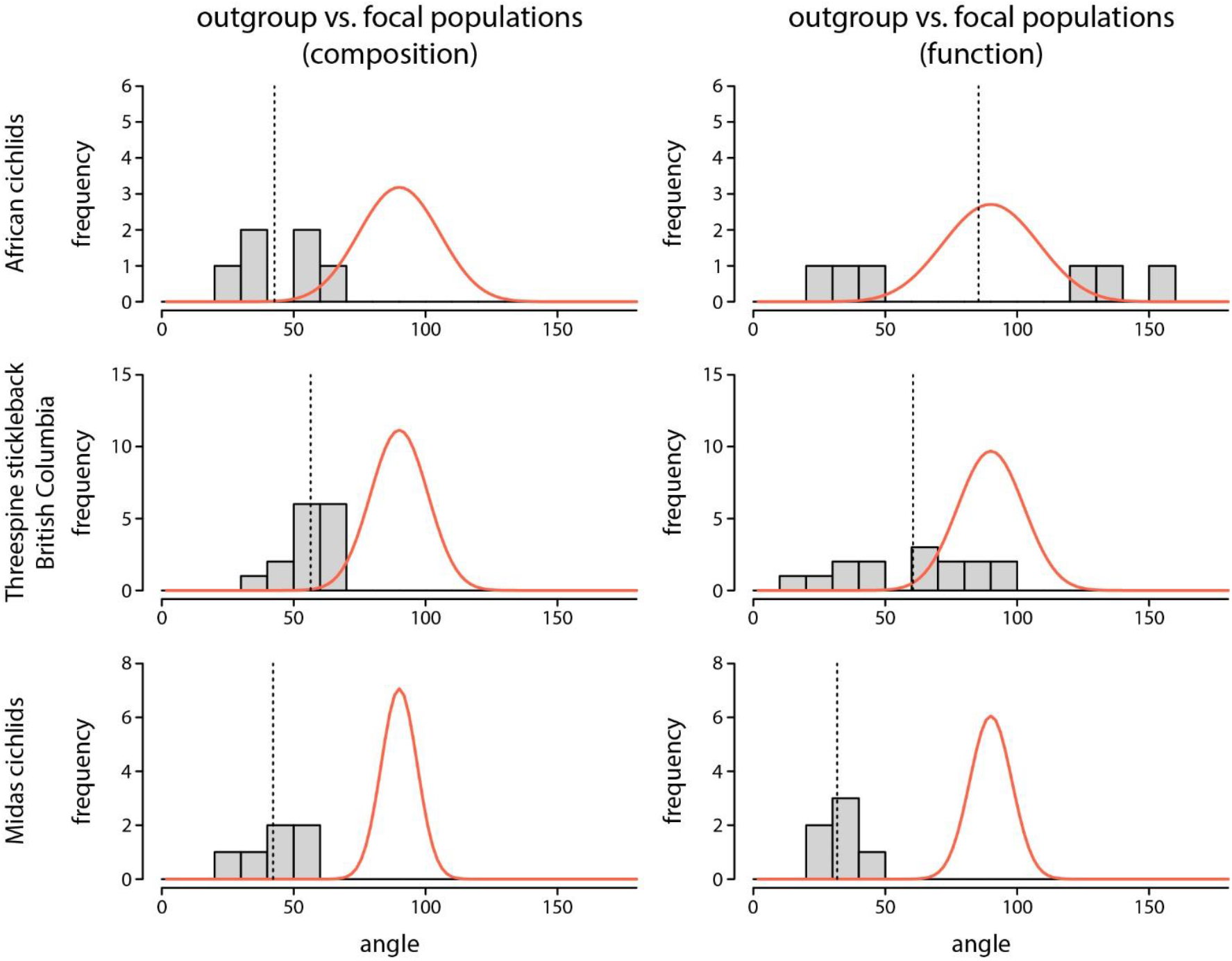
Histograms showing the distribution of angles between an outgroup and focal populations for gut microbiota composition (left column) and function (right column). The vertical dashed line represents the mean angle for each study system, and the multidimensional distribution of random angles is illustrated by the red curves, the shape of a curve is determined by the number of PCoA axes included in each analysis.

**Figure A7:**
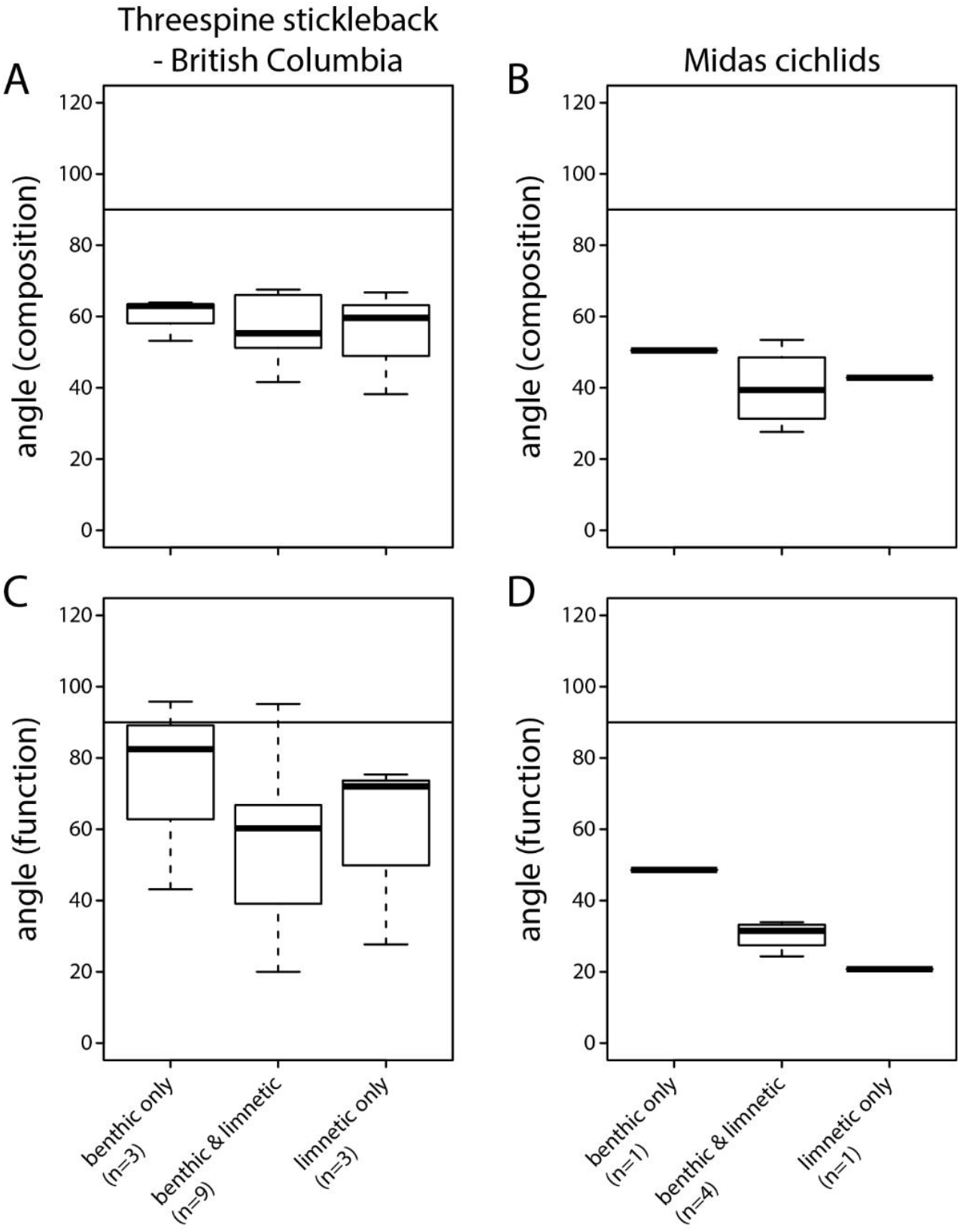
For the outgroup comparisons, we tested whether parallelism is stronger among focal populations of the same ecotype compared to focal populations of different ecotypes for gut microbiota composition and function in threespine stickleback from British Columbia (A&C) and Midas cichlids (B&D). The three different categories include angles of “outgroup-benthic vs. outgroup-benthic”, “outgroup-benthic vs. outgroup-limnetic” or “outgroup-limnetic vs. outgroup-limnetic” comparisons. We did not detect evidence that angles were smaller when only comparing among the same ecotype, suggesting that parallelism is mainly driven by the shared adaptation to a common environment among the focal populations, more so than by the adaptation to benthic and limnetic niches specifically.

**Table A1:**
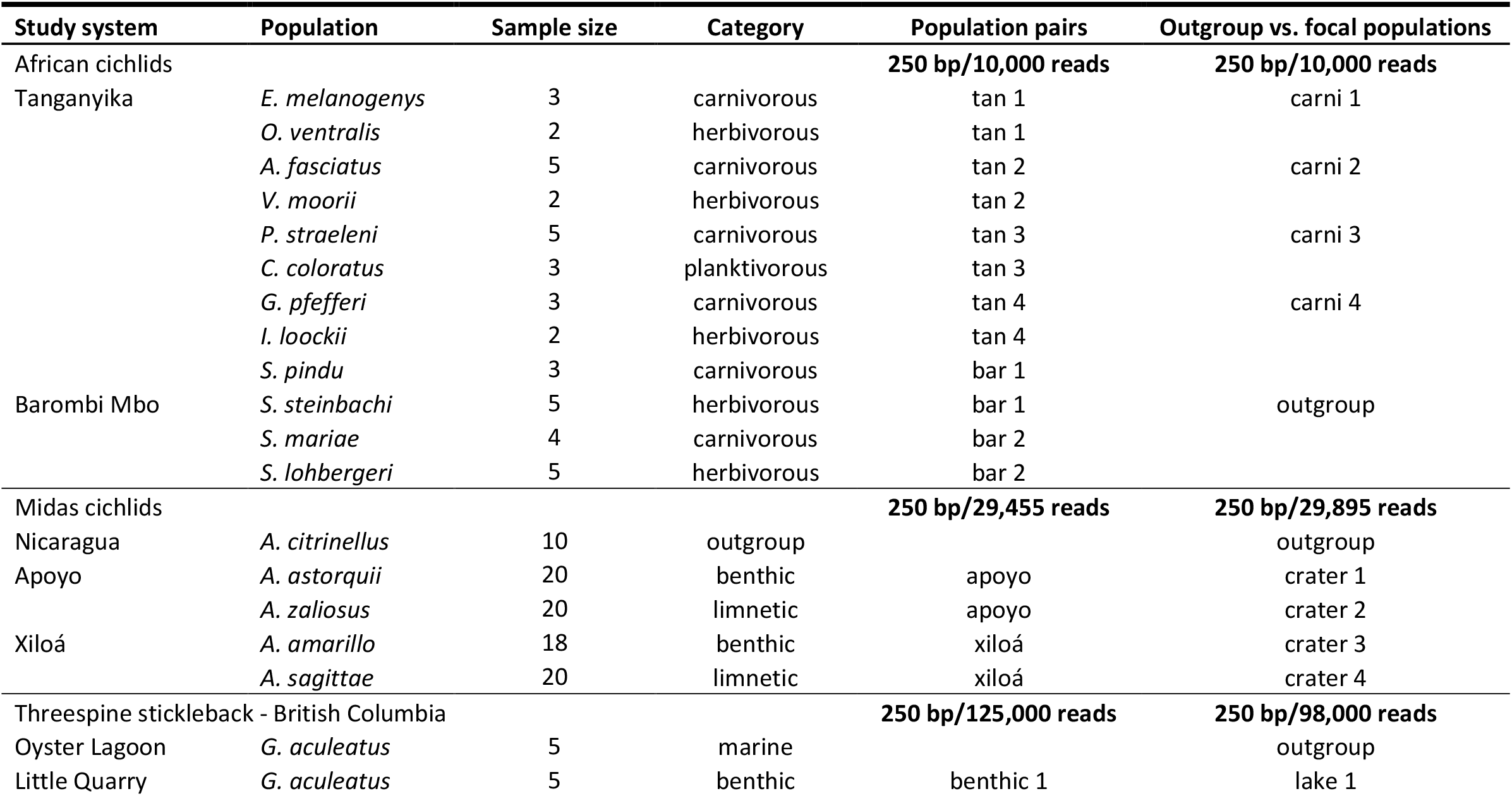
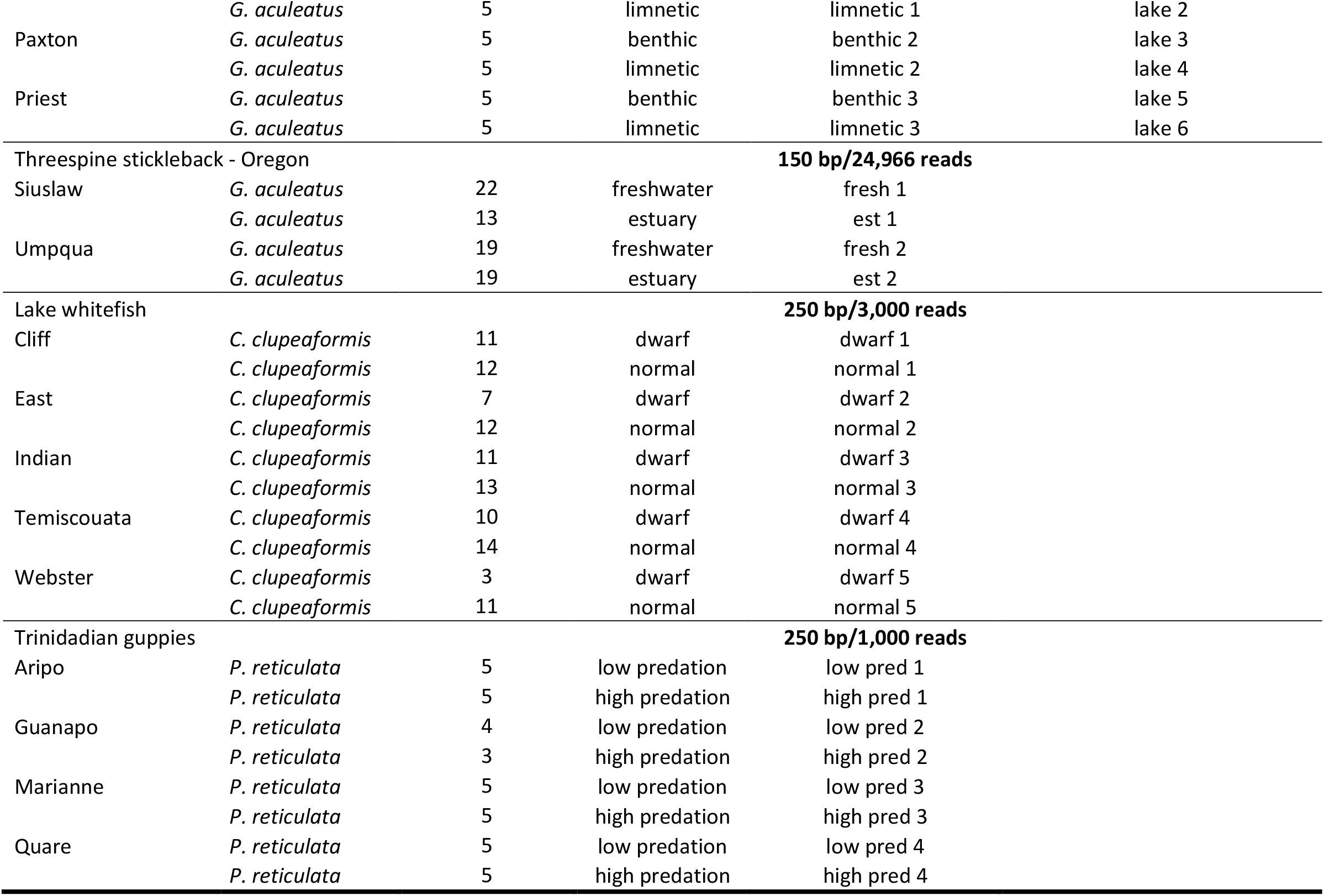
Identity of populations/species selected from each dataset that were included in our study, and information on their trophic ecology or habitat types as well as sample sizes. Different subsets were used in the population pair and outgroup vs. focal species analyses. For African cichlids, the herbivorous species from Barombi Mbo (*S. steinbachi*) represents an outgroup to the carnivorous species from Tanganyika based on a recent phylogeny (Irisarri et al., 2018). For threespine stickleback and Midas cichlids, outgroups are inferred ancestral populations that repeatedly colonized novel environments (Kautt et al., 2020, Bell and Foster, 1994). Sequencing read lengths and rarefaction depths used for all analyses are stated in bold for each study system and type of comparison.

**Table A2:**
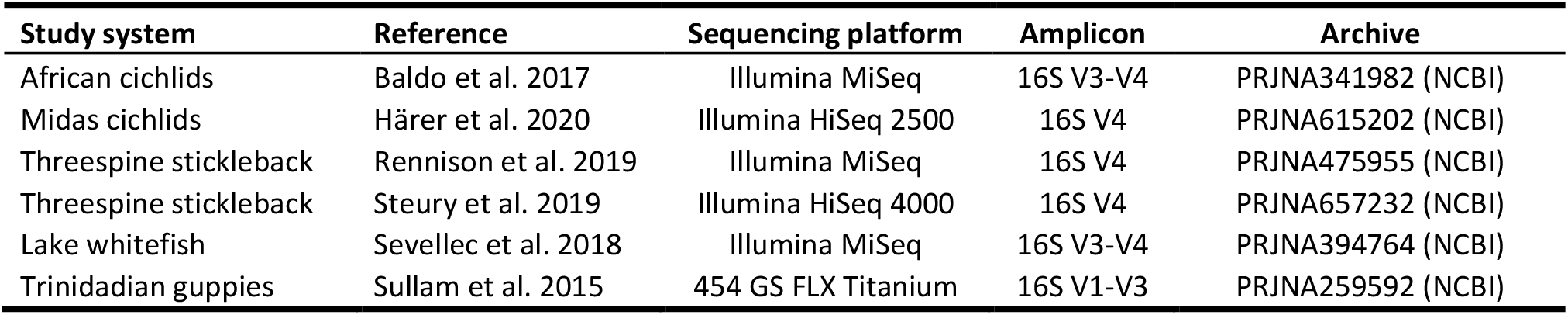
Information on sequencing platforms, amplified region and data archiving for all datasets included in this study.

| Study system | Reference | Sequencing platform | Amplicon | Archive |
| --- | --- | --- | --- | --- |
| African cichlids | Baldo et al. 2017 | Illumina MiSeq | 16S V3-V4 | PRJNA341982 (NCBI) |
| Midas cichlids | Härer et al. 2020 | Illumina HiSeq 2500 | 16S V4 | PRJNA615202 (NCBI) |
| Threespine stickleback | Rennison et al. 2019 | Illumina MiSeq | 16S V4 | PRJNA475955 (NCBI) |
| Threespine stickleback | Steury et al. 2019 | Illumina HiSeq 4000 | 16S V4 | PRJNA657232 (NCBI) |
| Lake whitefish | Sevellec et al. 2018 | Illumina MiSeq | 16S V3-V4 | PRJNA394764 (NCBI) |
| Trinidadian guppies | Sullam et al. 2015 | 454 GS FLX Titanium | 16S V1-V3 | PRJNA259592 (NCBI) |

**Table A3:** Inferred metagenome function was predicted with the PICRUSt2 plugin in QIIME2 with a maximum nearest-sequenced taxon index (NSTI) cutoff of 2, and reads above this value were discarded for these analyses. Across the study systems, a larger proportion of ASVs was below this cutoff (85.3-99.5%). Mean and median NSTI scores ranged from 0.45-2.689 and 0.072-0.366, respectively.

| Study | Mean(NSTI) | Median(NSTI) | Proportion ASVs > 2 |
| --- | --- | --- | --- |
| African cichlids (population pairs) | 0.465 | 0.366 | 0.005 |
| African cichlids (outgroup vs. focal populations) | 0.45 | 0.346 | 0.008 |
| Threespine stickleback - British Columbia (population pairs) | 0.853 | 0.061 | 0.031 |
| Threespine stickleback - British Columbia (outgroup vs. focal populations) | 0.532 | 0.102 | 0.028 |
| Trinidadian guppies (population pairs) | 0.606 | 0.072 | 0.06 |
| Lake whitefish (population pairs) | 2.689 | 0.078 | 0.147 |
| Threespine stickleback - Oregon (population pairs) | 1.231 | 0.332 | 0.073 |
| Midas cichlids (population pairs) | 0.939 | 0.156 | 0.039 |
| Midas cichlids (outgroup vs. focal populations) | 0.898 | 0.158 | 0.037 |

**Table A4:** Comparison of mean angles and statistical tests for parallelism (angles < 90°) for taxonomic composition of the gut microbiota across three different metrics: Bray-Curtis dissimilarity, weighted and unweighted UniFrac. Statistical significance was determined at the 0.05 level and results were highly consistent across metrics, only for threespine stickleback from British Columbia and lake whitefish did we detect some suggestive evidence (P < 0.1) for parallelism for one of the three metrics, but not in the other two. We performed one-sample t-tests when data was normally distributed; for data with non-normal distribution one-sample Wilcoxon signed-rank tests were used (indicated by asterisks). The dimensionality of each data set is indicated in brackets.

| **Taxonomic composition - Study system** | **Mean ($\gamma$)** | **P-value** |
| --- | --- | --- |
| ***African cichlids*** |  |  |
| Population pairs: Bray-Curtis (37) | 68.889 | **6.29E-06** |
| Population pairs: Unweighted UniFrac (41) | 66.245 | **5.94E-08** |
| Population pairs: Weighted UniFrac (28) | 57.158 | **7.09E-06** |
| Outgroup vs focal populations: Bray-Curtis (16) | 42.695 | **5.92E-04** |
| Outgroup vs focal populations: Unweighted UniFrac (16) | 50.373 | **3.34E-04** |
| Outgroup vs focal populations: Weighted UniFrac (13) | 60.667 | **0.023** |
| ***Threespine stickleback - British Columbia*** |  |  |
| Population pairs: Bray-Curtis* (26) | 81.016 | **0.087** |
| Population pairs: Unweighted UniFrac (28) | 87.471 | 0.119 |
| Population pairs: Weighted UniFrac* (21) | 80.844 | 0.625 |
| Outgroup vs focal populations: Bray-Curtis (30) | 56.421 | **1.15E-09** |
| Outgroup vs focal populations: Unweighted UniFrac (33) | 49.217 | **7.41E-15** |
| Outgroup vs focal populations: Weighted UniFrac (23) | 59.771 | **2.80E-07** |
| ***Lake whitefish*** |  |  |
| Population pairs: Bray-Curtis (80) | 90.132 | 0.511 |
| Population pairs: Unweighted UniFrac (98) | 85.056 | **0.072** |
| Population pairs: Weighted UniFrac (60) | 90.407 | 0.531 |
| ***Trinidadian guppies*** |  |  |
| Population pairs: Bray-Curtis (39) | 105.606 | 0.899 |
| Population pairs: Unweighted UniFrac (29) | 91.043 | 0.599 |
| Population pairs: Weighted UniFrac (21) | 87.682 | 0.366 |
| ***Midas cichlids*** |  |  |
| Outgroup vs focal populations: Bray-Curtis (73) | 42.179 | **3.26E-05** |
| Outgroup vs focal populations: Unweighted UniFrac (87) | 47.776 | **1.04E-05** |
| Outgroup vs focal populations: Weighted UniFrac (54) | 55.921 | **5.42E-04** |

**Table A5:** Comparison of mean angles and statistical tests for parallelism (angles < 90°) for inferred metagenome function across three different metrics based on Bray-Curtis dissimilarity: MetaCyc pathway abundances (PWAB), Enzyme Commission numbers (EC) and Kyoto Encyclopedia of Genes and Genomes orthologs (KO). Statistical significance was determined at the 0.05 level and results were consistent across metrics for all study systems. We performed one-sample t-tests when data was normally distributed; for data with non-normal distribution one-sample Wilcoxon signed-rank tests were used (indicated by asterisks). The dimensionality of each data set is indicated in brackets.

| Metagenome function - Study system | Mean ($\gamma$) | P-value |
| --- | --- | --- |
| ***African cichlids*** |  |  |
| Population pairs: PWAB (27) | 38.923 | **2.73E-09** |
| Population pairs: EC* (30) | 35.546 | **3.05E-05** |
| Population pairs: KO* (29) | 41.443 | **3.05E-05** |
| Outgroup vs focal populations: PWAB (12) | 85.38 | 0.425 |
| Outgroup vs focal populations: EC (12) | 80.94 | 0.341 |
| Outgroup vs focal populations: KO (11) | 82.325 | 0.351 |
| ***Threespine stickleback - British Columbia*** |  |  |
| Population pairs: PWAB (21) | 88.045 | 0.471 |
| Population pairs: EC (20) | 89.19 | 0.488 |
| Population pairs: KO (20) | 84.645 | 0.413 |
| Outgroup vs focal populations: PWAB (23) | 60.485 | **1.93E-04** |
| Outgroup vs focal populations: EC (29) | 54.664 | **4.77E-06** |
| Outgroup vs focal populations: KO (22) | 53.186 | **2.36E-06** |
| ***Lake whitefish*** |  |  |
| Population pairs: PWAB (65) | 91.112 | 0.566 |
| Population pairs: EC (68) | 90.883 | 0.552 |
| Population pairs: KO (64) | 91.456 | 0.584 |
| ***Trinidadian guppies*** |  |  |
| Population pairs: PWAB (28) | 95.56 | 0.69 |
| Population pairs: EC (28) | 94.729 | 0.716 |
| Population pairs: KO (28) | 97.021 | 0.766 |
| ***Midas cichlids*** |  |  |
| Outgroup vs focal populations: PWAB (54) | 31.761 | **1.29E-05** |
| Outgroup vs focal populations: EC (57) | 32.745 | **2.20E-05** |
| Outgroup vs focal populations: KO (55) | 28.123 | **7.62E-06** |

## Notes

### Competing Interest Statement

The authors have declared no competing interest.

## References

Adams, D. C. & Collyer, M. L. 2009. A general framework for the analysis of phenotypic trajectories in evolutionary studies. Evolution, 63, 1143–54.

Amato, K. R., Sanders, J. G., Song, S. J., Nute, M., Metcalf, J. L., Thompson, L. R., Morton, J. T., Amir, A., V, J. M., Humphrey, G., Gogul, G., Gaffney, J., A, L. B., G, A. O. B., F, P. C., Di Fiore, A., N, J. D., T, L. G., Gomez, A., Kowalewski, M. M., R, J. L., Link, A., M, L. S., Tecot, S., B, A. W., K, E. N., R, M. S., Knight, R. & S, R. L. 2019. Evolutionary trends in host physiology outweigh dietary niche in structuring primate gut microbiomes. The ISME Journal, 13, 576–587.

Baldo, L., Pretus, J. L., Riera, J. L., Musilova, Z., Bitja Nyom, A. R. & Salzburger, W. 2017. Convergence of gut microbiotas in the adaptive radiations of African cichlid fishes. The ISME Journal, 11, 1975–1987.

Baniel, A., Amato, K. R., Beehner, J. C., Bergman, T. J., Mercer, A., Perlman, R. F., Petrullo, L., Reitsema, L., Sams, S., Lu, A. & Snyder-Mackler, N. 2021. Seasonal shifts in the gut microbiome indicate plastic responses to diet in wild geladas. Microbiome, 9, 26.

Barluenga, M. & Meyer, A. 2010. Phylogeography, colonization and population history of the Midas cichlid species complex (Amphilophus spp.) in the Nicaraguan crater lakes. BMC Evolutionary Biology, 10, 326.

Bell, M. A. & Foster, S. A. 1994. The evolutionary biology of the threespine stickleback. Oxford University Press, Oxford.

Benson, A. K., Kelly, S. A., Legge, R., Ma, F. R., Low, S. J., Kim, J., Zhang, M., Oh, P. L., Nehrenberg, D., Hua, K. J., Kachman, S. D., Moriyama, E. N., Walter, J., Peterson, D. A. & Pomp, D. 2010. Individuality in gut microbiota composition is a complex polygenic trait shaped by multiple environmental and host genetic factors. Proceedings of the National Academy of Sciences of the United States of America, 107, 18933–18938.

Bernatchez, L., Chouinard, A. & Lu, G. Q. 1999. Integrating molecular genetics and ecology in studies of adaptive radiation: whitefish, Coregonus sp., as a case study. Biological Journal of the Linnean Society, 68, 173–194.

Bletz, M. C., Goedbloed, D. J., Sanchez, E., Reinhardt, T., Tebbe, C. C., Bhuju, S., Geffers, R., Jarek, M., Vences, M. & Steinfartz, S. 2016. Amphibian gut microbiota shifts differentially in community structure but converges on habitat-specific predicted functions. Nature Communications, 7, 13699.

Bolnick, D. I., Barrett, R. D. H., Oke, K. B., Rennison, D. J. & Stuart, Y. E. 2018. (Non)parallel evolution. Annual Review of Ecology, Evolution, and Systematics, 49, 303–330.

Bolnick, D. I., Snowberg, L. K., Hirsch, P. E., Lauber, C. L., Knight, R., Caporaso, J. G. & Svanback, R. 2014a. Individuals’ diet diversity influences gut microbial diversity in two freshwater fish (threespine stickleback and Eurasian perch). Ecology Letters, 17, 979–987.

Bolnick, D. I., Snowberg, L. K., Hirsch, P. E., Lauber, C. L., Org, E., Parks, B., Lusis, A. J., Knight, R., Caporaso, J. G. & Svanback, R. 2014b. Individual diet has sex-dependent effects on vertebrate gut microbiota. Nature Communications, 5, 4500.

Bolyen, E., Rideout, J. R., Dillon, M. R., Bokulich, N. A., Abnet, C. C., Al-Ghalith, G. A., Alexander, H., Alm, E. J., Arumugam, M., Asnicar, F., Bai, Y., Bisanz, J. E., Bittinger, K., Brejnrod, A., Brislawn, C. J., Brown, C. T., Callahan, B. J., Caraballo-Rodriguez, A. M., Chase, J., Cope, E. K., Da Silva, R., Diener, C., Dorrestein, P. C., Douglas, G. M., Durall, D. M., Duvallet, C., Edwardson, C. F., Ernst, M., Estaki, M., Fouquier, J., Gauglitz, J. M., Gibbons, S. M., Gibson, D. L., Gonzalez, A., Gorlick, K., Guo, J., Hillmann, B., Holmes, S., Holste, H., Huttenhower, C., Huttley, G. A., Janssen, S., Jarmusch, A. K., Jiang, L., Kaehler, B. D., Kang, K. B., Keefe, C. R., Keim, P., Kelley, S. T., Knights, D., Koester, I., Kosciolek, T., Kreps, J., Langille, M. G. I., Lee, J., Ley, R., Liu, Y. X., Loftfield, E., Lozupone, C., Maher, M., Marotz, C., Martin, B. D., Mcdonald, D., Mciver, L. J., Melnik, A. V., Metcalf, J. L., Morgan, S. C., Morton, J. T., Naimey, A. T., Navas-Molina, J. A., Nothias, L. F., Orchanian, S. B., Pearson, T., Peoples, S. L., Petras, D., Preuss, M. L., Pruesse, E., Rasmussen, L. B., Rivers, A., Robeson, M. S., 2nd, Rosenthal, P., Segata, N., Shaffer, M., Shiffer, A., Sinha, R., Song, S. J., Spear, J. R., Swafford, A. D., Thompson, L. R., Torres, P. J., Trinh, P., Tripathi, A., Turnbaugh, P. J., Ul-Hasan, S., Van Der Hooft, J. J. J., Vargas, F., Vazquez-Baeza, Y., Vogtmann, E., Von Hippel, M., Walters, W., et al. 2019. Reproducible, interactive, scalable and extensible microbiome data science using QIIME 2. Nature Biotechnology, 37, 852–857.

Brooks, A. W., Kohl, K. D., Brucker, R. M., Van Opstal, E. J. & Bordenstein, S. R. 2016. Phylosymbiosis: relationships and functional effects of microbial communities across host evolutionary history. PLoS Biology, 14, e2000225.

Burns, A. R., Miller, E., Agarwal, M., Rolig, A. S., Milligan-Myhre, K., Seredick, S., Guillemin, K. & Bohannan, B. J. M. 2017. Interhost dispersal alters microbiome assembly and can overwhelm host innate immunity in an experimental zebrafish model. Proceedings of the National Academy of Sciences of the United States of America, 114, 11181–11186.

Callahan, B. J., Mcmurdie, P. J., Rosen, M. J., Han, A. W., Johnson, A. J. A. & Holmes, S. P. 2016. DADA2: High-resolution sample inference from Illumina amplicon data. Nature Methods, 13, 581–583.

Collyer, M. L. & Adams, D. C. 2007. Analysis of two-state multivariate phenotypic change in ecological studies. Ecology, 88, 683–92.

Colosimo, P. F., Hosemann, K. E., Balabhadra, S., Villarreal, G., Jr., Dickson, M., Grimwood, J., Schmutz, J., Myers, R. M., Schluter, D. & Kingsley, D. M. 2005. Widespread parallel evolution in sticklebacks by repeated fixation of Ectodysplasin alleles. Science, 307, 1928–33.

Costello, E. K., Stagaman, K., Dethlefsen, L., Bohannan, B. J. M. & Relman, D. A. 2012. The application of ecological theory toward an understanding of the human microbiome. Science, 336, 1255–1262.

Delsuc, F., Metcalf, J. L., Wegener Parfrey, L., Song, S. J., Gonzalez, A. & Knight, R. 2014. Convergence of gut microbiomes in myrmecophagous mammals. Molecular Ecology, 23, 1301–17.

Douglas, G. M., Maffei, V. J., Zaneveld, J. R., Yurgel, S. N., Brown, J. R., Taylor, C. M., Huttenhower, C. & Langille, M. G. I. 2020. PICRUSt2 for prediction of metagenome functions. Nat Biotechnol, 38, 685–688.

Elmer, K. R., Fan, S., Kusche, H., Spreitzer, M. L., Kautt, A. F., Franchini, P. & Meyer, A. 2014. Parallel evolution of Nicaraguan crater lake cichlid fishes via non-parallel routes. Nature Communications, 5, 5168.

Elmer, K. R., Kusche, H., Lehtonen, T. K. & Meyer, A. 2010. Local variation and parallel evolution: morphological and genetic diversity across a species complex of neotropical crater lake cichlid fishes. Philosophical Transactions of the Royal Society B-Biological Sciences, 365, 1763–82.

Goodrich, J. K., Waters, J. L., Poole, A. C., Sutter, J. L., Koren, O., Blekhman, R., Beaumont, M., Van Treuren, W., Knight, R., Bell, J. T., Spector, T. D., Clark, A. G. & Ley, R. E. 2014. Human genetics shape the gut microbiome. Cell, 159, 789–799.

Härer, A., Torres-Dowdall, J., Rometsch, S. J., Yohannes, E., Machado-Schiaffino, G. & Meyer, A. 2020. Parallel and non-parallel changes of the gut microbiota during trophic diversification in repeated young adaptive radiations of sympatric cichlid fish. Microbiome, 8, 149.

Human Microbiome Project Consortium 2012. Structure, function and diversity of the healthy human microbiome. Nature, 486, 207–14.

Irisarri, I., Singh, P., Koblmüller, S., Torres-Dowdall, J., Henning, F., Franchini, P., Fischer, C., Lemmon, A., Lemmon, E., Thallinger, G., Sturmbauer, C. & Meyer, A. 2018. Phylogenomics uncovers early hybridization and adaptive loci shaping the radiation of Lake Tanganyika cichlid fishes. Nature Communications, 9, 3159.

Kanehisa, M., Goto, S., Sato, Y., Furumichi, M. & Tanabe, M. 2012. KEGG for integration and interpretation of large-scale molecular data sets. Nucleic Acids Research, 40, D109–14.

Kautt, A. F., Kratochwil, C. F., Nater, A., Machado-Schiaffino, G., Olave, M., Henning, F., Torres-Dowdall, J., Härer, A., Hulsey, C. D., Franchini, P., Pippel, M., Myers, E. W. & Meyer, A. 2020. Contrasting signatures of genomic divergence during sympatric speciation. Nature, 588, 106–111.

Kolodny, O. & Schulenburg, H. 2020. Microbiome-mediated plasticity directs host evolution along several distinct time scales. Philosophical Transactions of the Royal Society B-Biological Sciences, 375, 20190589.

Ley, R., Hamady, M., Lozupone, C., Turnbaugh, P. J., Ramey, R. R., Bircher, J. S., Schlegel, M. L., Tucker, T. A., Schrenzel, M. D., Knight, R. & Gordon, J. I. 2008a. Evolution of mammals and their gut microbes. Science, 320, 1647–1651.

Ley, R., Lozupone, C., Hamady, M., Knight, R. & Gordon, J. I. 2008b. Worlds within worlds: evolution of the vertebrate gut microbiota. Nature Reviews Microbiology, 6, 776–788.

Ley, R. E., Peterson, D. A. & Gordon, J. I. 2006. Ecological and evolutionary forces shaping microbial diversity in the human intestine. Cell, 124, 837–848.

Li, T., Long, M., Li, H., Gatesoupe, F. J., Zhang, X., Zhang, Q., Feng, D. & Li, A. 2017. Multi-omics analysis reveals a correlation between the host phylogeny, gut microbiota and metabolite profiles in cyprinid fishes. Frontiers in Microbiology, 8, doi: 10.3389/fmicb.2017.00454.

Losos, J. B., Jackman, T. R., Larson, A., Queiroz, K. & Rodriguez-Schettino, L. 1998. Contingency and determinism in replicated adaptive radiations of island lizards. Science, 279, 2115–8.

Lozupone, C., Lladser, M. E., Knights, D., Stombaugh, J. & Knight, R. 2011. UniFrac: an effective distance metric for microbial community comparison. The ISME Journal, 5, 169–72.

Mardia, K., Kent, J. & Bibby, J. 1979. Multivariate analysis. Academic Press, London.

Martinez, I., Maldonado-Gomez, M. X., Gomes-Neto, J. C., Kittana, H., Ding, H., Schmaltz, R., Joglekar, P., Cardona, R. J., Marsteller, N. L., Kembel, S. W., Benson, A. K., Peterson, D. A., Ramer-Tait, A. E. & Walter, J. 2018. Experimental evaluation of the importance of colonization history in early-life gut microbiota assembly. eLife, 7, e36521.

Matthews, B., Marchinko, K. B., Bolnick, D. I. & Mazumder, A. 2010. Specialization of trophic position and habitat use by sticklebacks in an adaptive radiation. Ecology, 91, 1025–1034.

Muegge, B. D., Kuczynski, J., Knights, D., Clemente, J. C., Gonzalez, A., Fontana, L., Henrissat, B., Knight, R. & Gordon, J. I. 2011. Diet Drives Convergence in Gut Microbiome Functions Across Mammalian Phylogeny and Within Humans. Science, 332, 970–974.

Mulder, I. E., Schmidt, B., Stokes, C. R., Lewis, M., Bailey, M., Aminov, R. I., Prosser, J. I., Gill, B. P., Pluske, J. R., Mayer, C. D., Musk, C. C. & Kelly, D. 2009. Environmentally-acquired bacteria influence microbial diversity and natural innate immune responses at gut surfaces. BMC Biology, 7, 79.

Muschick, M., Indermaur, A. & Salzburger, W. 2012. Convergent Evolution within an Adaptive Radiation of Cichlid Fishes. Current Biology, 22, 2362–2368.

Ormond, C. I., Rosenfeld, J. S. & Taylor, E. B. 2011. Environmental determinants of threespine stickleback species pair evolution and persistence. Canadian Journal of Fisheries and Aquatic Sciences, 68, 1983–1997.

Pedregosa, F., Varoquaux, G., Gramfort, A., Michel, V., Thirion, B., Grisel, O., Blondel, M., Prettenhofer, P., Weiss, R., Dubourg, V., Vanderplas, J., Passos, A., Cournapeau, D., Brucher, M., Perrot, M. & Duchesnay, E. 2011. Scikit-learn: Machine Learning in Python. Journal of Machine Learning Research, 12, 2825–2830.

Price, M. N., Dehal, P. S. & Arkin, A. P. 2010. FastTree 2-approximately maximum-likelihood trees for large alignments. PLoS One, 5, e9490.

Quast, C., Pruesse, E., Yilmaz, P., Gerken, J., Schweer, T., Yarza, P., Peplies, J. & Glockner, F. O. 2013. The SILVA ribosomal RNA gene database project: improved data processing and web-based tools. Nucleic Acids Research, 41, 590–596.

R_Core_Team 2021. R: A language and environment for statistical computing, Vienna, Austria. Retrieved from https://www.R-project.org/.

Rennison, D. J., Rudman, S. M. & Schluter, D. 2019. Parallel changes in gut microbiome composition and function during colonization, local adaptation and ecological speciation. Proceedings of the Royal Society B-Biological Sciences, 286, 20191911.

Reznick, D., Rodd, F. H. & Cardenas, M. 1996. Life-history evolution in guppies (Poecilia reticulata: Poeciliidae). IV. Parallelism in life-history phenotypes. American Naturalist, 147, 319–338.

Rosenblum, E. B., Parent, C. E., Diepeveen, E. T., Noss, C. & Bi, K. 2017. Convergent phenotypic evolution despite contrasting demographic histories in the fauna of White Sands. American Naturalist, 190, S45–S56.

Rudman, S. M., Greenblum, S., Hughes, R. C., Rajpurohit, S., Kiratli, O., Lowder, D. B., Lemmon, S. G., Petrov, D. A., Chaston, J. M. & Schmidt, P. 2019. Microbiome composition shapes rapid genomic adaptation of Drosophila melanogaster. Proceedings of the National Academy of Sciences of the United States of America, 116, 20025–20032.

Schluter, D. & Mcphail, J. D. 1992. Ecological character displacement and speciation in sticklebacks. American Naturalist, 140, 85–108.

Sevellec, M., Derome, N. & Bernatchez, L. 2018. Holobionts and ecological speciation: the intestinal microbiota of lake whitefish species pairs. Microbiome, 6, 47.

Shapiro, S. S. & Wilk, M. B. 1965. An analysis of variance test for normality (complete samples). Biometrika, 52, 591–611.

Smith, C. C. R., Snowberg, L. K., Caporaso, J. G., Knight, R. & Bolnick, D. I. 2015. Dietary input of microbes and host genetic variation shape among-population differences in stickleback gut microbiota. The ISME Journal, 9, 2515–2526.

Smits, S. A., Leach, J., Sonnenburg, E. D., Gonzalez, C. G., Lichtman, J. S., Reid, G., Knight, R., Manjurano, A., Changalucha, J., Elias, J. E., Dominguez-Bello, M. G. & Sonnenburg, J. L. 2017. Seasonal cycling in the gut microbiome of the Hadza hunter-gatherers of Tanzania. Science, 357, 802–806.

Song, S. J., Sanders, J. G., Delsuc, F., Metcalf, J., Amato, K., Taylor, M. W., Mazel, F., Lutz, H. L., Winker, K., Graves, G. R., Humphrey, G., Gilbert, J. A., Hackett, S. J., White, K. P., Skeen, H. R., Kurtis, S. M., Withrow, J., Braile, T., Miller, M., Mccracken, K. G., Maley, J. M., Ezenwa, V. O., Williams, A., Blanton, J. M., Mckenzie, V. J. & Knight, R. 2020. Comparative analyses of vertebrate gut microbiomes reveal convergence between birds and bats. mBio, 11, e02901–19.

Spor, A., Koren, O. & Ley, R. 2011. Unravelling the effects of the environment and host genotype on the gut microbiome. Nature Reviews Microbiology, 9, 279–290.

Steiner, C. C., Rompler, H., Boettger, L. M., Schoneberg, T. & Hoekstra, H. E. 2009. The genetic basis of phenotypic convergence in Beach Mice: Similar pigment patterns but different genes. Molecular Biology and Evolution, 26, 35–45.

Steury, R. A., Currey, M. C., Cresko, W. A. & Bohannan, B. J. M. 2019. Population genetic divergence and environment influence the gut microbiome in Oregon threespine stickleback. Genes, 10, 484.

Stuart, Y. E., Veen, T., Weber, J. N., Hanson, D., Ravinet, M., Lohman, B. K., Thompson, C. J., Tasneem, T., Doggett, A., Izen, R., Ahmed, N., Barrett, R. D. H., Hendry, A. P., Peichel, C. L. & Bolnick, D. I. 2017. Contrasting effects of environment and genetics generate a continuum of parallel evolution. Nature Ecology & Evolution, 1, 0158.

Sullam, K. E., Essinger, S. D., Lozupone, C. A., O’connor, M. P., Rosen, G. L., Knight, R., Kilham, S. S. & Russell, J. A. 2012. Environmental and ecological factors that shape the gut bacterial communities of fish: a meta-analysis. Molecular Ecology, 21, 3363–3378.

Sullam, K. E., Rubin, B. E., Dalton, C. M., Kilham, S. S., Flecker, A. S. & Russell, J. A. 2015. Divergence across diet, time and populations rules out parallel evolution in the gut microbiomes of Trinidadian guppies. The ISME Journal, 9, 1508–22.

Sun, S., Jones, R. B. & Fodor, A. A. 2020. Inference-based accuracy of metagenome prediction tools varies across sample types and functional categories. Microbiome, 8, 46.

Tarnecki, A. M., Burgos, F. A., Ray, C. L. & Arias, C. R. 2017. Fish intestinal microbiome: diversity and symbiosis unravelled by metagenomics. Journal of Applied Microbiology, 123, 2–17.

Taylor, E. B. & Mcphail, J. D. 1999. Evolutionary history of an adaptive radiation in species pairs of threespine sticklebacks (Gasterosteus): insights from mitochondrial DNA. Biological Journal of the Linnean Society, 66, 271–291.

Thompson, K. A., Osmond, M. M. & Schluter, D. 2019. Parallel genetic evolution and speciation from standing variation. Evolution Letters, 3, 129–141.

Torres-Dowdall, J., Pierotti, M. E. R., Härer, A., Karagic, N., Woltering, J. M., Henning, F., Elmer, K. R. & Meyer, A. 2017. Rapid and parallel adaptive evolution of the visual system of Neotropical Midas cichlid fishes. Molecular Biology and Evolution, 34, 2469–2485.

Turnbaugh, P. J., Ley, R. E., Mahowald, M. A., Magrini, V., Mardis, E. R. & Gordon, J. I. 2006. An obesity-associated gut microbiome with increased capacity for energy harvest. Nature, 444, 1027–1031.

Turnbaugh, P. J., Ridaura, V. K., Faith, J. J., Rey, F. E., Knight, R. & Gordon, J. I. 2009. The effect of diet on the human gut microbiome: a metagenomic analysis in humanized gnotobiotic mice. Science Translational Medicine, 1, 6ra14.

Watanabe, J. 2022. Detecting (non)parallel evolution in multidimensional spaces: angles, correlations and eigenanalysis. Biol Lett, 18, 20210638.

Wilcoxon, F. 1945. Individual comparisons by ranking methods. Biometrics Bulletin, 1, 80–83.

Youngblut, N. D., Reischer, G. H., Walters, W., Schuster, N., Walzer, C., Stalder, G., Ley, R. E. & Farnleitner, A. H. 2019. Host diet and evolutionary history explain different aspects of gut microbiome diversity among vertebrate clades. Nature Communications, 10, 2200.

Zepeda Mendoza, M. L., Xiong, Z., Escalera-Zamudio, M., Runge, A. K., Theze, J., Streicker, D., Frank, H. K., Loza-Rubio, E., Liu, S., Ryder, O. A., Samaniego Castruita, J. A., Katzourakis, A., Pacheco, G., Taboada, B., Lober, U., Pybus, O. G., Li, Y., Rojas-Anaya, E., Bohmann, K., Carmona Baez, A., Arias, C. F., Liu, S., Greenwood, A. D., Bertelsen, M. F., White, N. E., Bunce, M., Zhang, G., Sicheritz-Ponten, T. & Gilbert, M. P. T. 2018. Hologenomic adaptations underlying the evolution of sanguivory in the common vampire bat. Nature Ecology & Evolution, 2, 659–668.

Zhang, M. L., Sun, Y. H., Liu, Y. K., Qiao, F., Chen, L. Q., Liu, W. T., Du, Z. Y. & Li, E. C. 2016. Response of gut microbiota to salinity change in two euryhaline aquatic animals with reverse salinity preference. Aquaculture, 454, 72–80.

